# Systematic annotation of *Helitron*-like elements in eukaryote genomes using HELIANO

**DOI:** 10.1101/2024.02.08.579435

**Authors:** Zhen Li, Clément Gilbert, Haoran Peng, Nicolas Pollet

## Abstract

Helitron-like elements (HLEs) are widespread eukaryotic DNA transposons employing a rolling-circle transposition mechanism. Despite their prevalence in fungi, animals, and plant genomes, identifying *Helitrons* remains challenging. We introduce HELIANO, a software for annotating and classifying autonomous and non-autonomous *Helitron* and *Helentron* sequences from whole genomes. HELIANO outperforms existing tools in speed and accuracy, demonstrated through benchmarking and its application to complex genomes (*Xenopus tropicalis, Xenopus laevis, Oryza sativa*), revealing numerous newly identified *Helitrons* and *Helentrons*.

In a comprehensive analysis of 404 eukaryote genomes, we found HLEs widely distributed across phyla, with exceptions in specific taxa. *Helentrons* were identified in numerous land plant species, and 20 protein domains were discovered integrated within specific autonomous HLE families. A global phylogenetic analysis confirmed the classification into main clades *Helentron* and *Helitron*, revealing nine subgroups, some enriched in particular taxa. The future use of HELIANO will contribute to the global analysis of TEs across genomes and enhance our understanding of this transposon superfamily.

## Introduction

Transposable elements (TEs) are ubiquitous selfish genetic elements characterized by their capacity to move and duplicate within genomes (1, 2). The nature and abundance of TEs exhibit substantial variation across species (3, 4). This diversity of the TE landscape across genomes is associated with the absence of an ideal TE prediction tool, and manual curation will likely remain the best way to obtain the most accurate map of TEs in a given genome (5, 6). Yet, as complete genome sequences from all over the Tree of Life are produced, our global understanding of TEs diversity, evolution and impact on host genomes will continue to benefit strongly from improving automated algorithms that enable fast TE identification and classification (1, 4, 6, 7). Here, we present a new algorithm dedicated to fast and accurate *de novo* annotation of *Helitrons* while overcoming several limitations of existing tools (8–10).

Among eukaryote DNA transposons, *Helitrons* form a particular superfamily believed to use a rolling-circle transposition mechanism to spread in genomes (11). Recent studies have identified variants of *Helitrons* named *Helentron* and *Helitron2* (12–17). To avoid confusion, we will collectively use the term *Helitron*-like elements (HLEs) to refer to all variants (Figure 1). HLEs have been reported in the genomes of numerous eukaryotic taxa, including fungi, animals, plants, and algae (8, 11, 12, 15, 17–20). Among some well-studied species, they can contribute a considerable proportion of their genome sequence (20, 21). For example, *Helitrons* have been estimated to span around 6% of the little brown bat genome and about 4% of the silkworm genome (20, 21). Numerous reports revealed that *Helitrons* can capture genes and lead to horizontal transfer and genome shuffling, making them significant sources for genome dynamics and evolution (18, 22–24). While their evolutionary significance is undebated, HLEs are still tricky to identify efficiently because they do not create target site duplication (TSD) upon transposition and lack classical structural features (1, 8).

**Figure 1.**
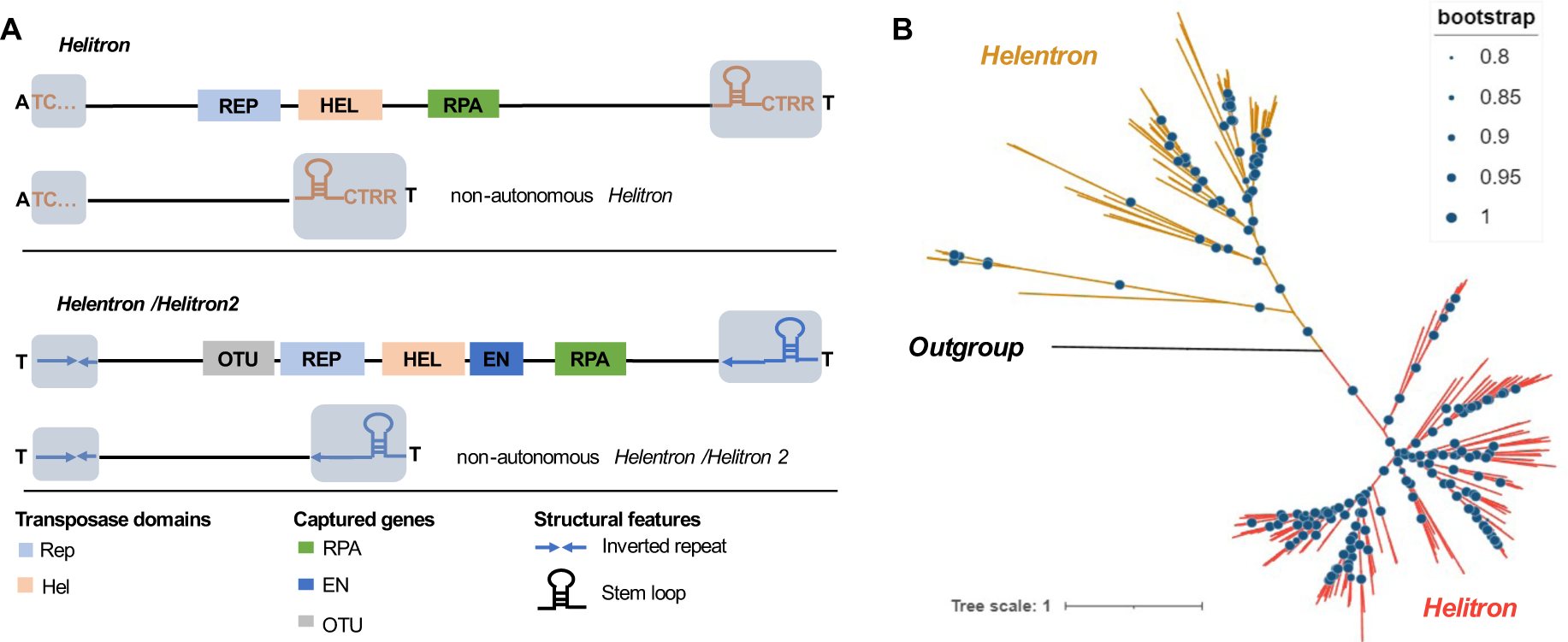
*Helitron*-Like Elements (HLEs) variants. (A) Features of HLEs. Autonomous elements are pictured on top, with their non-autonomous derivatives just below. Non-autonomous HLEs share identical Left Terminal Sequences (LTS) and Right Terminal Sequences (RTS) with their autonomous counterparts. All autonomous HLEs encode the Rep (light blue) and Helicase (orange) domains. *Helitrons* might also carry the RPA domain (green), and *Helentrons* might have the EN domain (blue), RPA domain, and OTU domain (grey). Variant *Helitrons* usually insert between A and T nucleotides, while *Helentrons* usually insert between T and T nucleotides. The scale of this scheme is relative. (B) Maximum likelihood estimation tree of HLE transposases from Repbase (LogLk = -153572.880) (27). The clade highlighted in red corresponds to the *Helitrons* variant, and the clade highlighted in orange corresponds to the *Helentrons* variant (including *Helitron 2*, see text). As an outgroup, we used a sequence made by concatenating geminivirus catalytic rep and helicase proteins of *Myroides phaeus*. Blue dots on the branches of the tree are bootstrap values greater than 0.8.

The HLE world includes three recurrent variants named *Helitron*, *Helentron*, and *Helitron2* (12). Although all autonomous HLEs encode Rep/Helicase (RepHel) transposase having both the Rep endonuclease and superfamily I helicase protein domains, additional characteristics enable their discrimination (Figure 1A, B) (12). In their terminal regions, *Helitrons* usually start with the TC signal and terminate with a hairpin structure with a suffix of CTRR motif (Figure 1A) (8, 11, 12). HLEs also vary in their insertion site preferences: the *Helitron* variant usually inserts between A and T nucleotides, while *Helitron2*/*Helentrons* generally insert between T and T nucleotides (12, 15, 16) (Figure 1A). Based on their transposase sequence similarity, *Helitron* and *Helentron/Helitron2* variants are phylogenetically distinguishable (12, 13, 25). Although some studies argued that *Helentron* and *Helitron2* are distinct variants, a more recent study based on a detailed phylogenetic analysis of RepHel sequences showed that the *Helitron2* variant is both phylogenetically and structurally undistinguishable from the *Helentron* variant (12, 13, 15). Thus, we use *Helentron* to refer to *Helentron* and *Helitron2* in this work.

Besides the RepHel transposase domain, many additional gene sequences are recurrently found in HLEs (12). For example, the gene encoding a single-stranded DNA-binding protein homologous to the replication protein A (RPA) can be detected in *Helitrons* and *Helentrons*. Similarly, the Ovarian Tumor protein (OTU) and apurinic/apyrimidinic (AP) endonuclease (EN) gene fragments could only be found in *Helentrons* (11, 12, 16, 17). These gene sequences tend to be fragmented and are not always detected in autonomous HLEs (12). Finally, autonomous HLEs often give birth to thousands of non-autonomous insertions, which share high similarity with their autonomous counterparts at both terminal regions (12) (Figure 1A).

Currently, tools for detecting HLEs are mainly structure-based, e.g., HelitronScanner, EAHelitron, and HelSearch (8, 10, 26). The primary strategy used in these tools is to search terminal signals of canonical *Helitrons*: the TC signals for the left terminal region and the CTRR motif for the right terminal region (20). Therefore, these tools cannot detect *Helentrons* or distinguish *Helentrons* from *Helitrons* (12). Furthermore, since such terminal signals are widespread in genome sequences, these software tools suffer from a high rate of false positives (8).

Our new software tool, HELIANO, a *Helitron*-like element annotator, was designed to comprehensively annotate all autonomous HLEs and their associated non-autonomous elements in a given genome. Unlike previously developed tools used for HLE identification, HELIANO first relies on homology-based searches for detecting autonomous HLEs and then characterizes candidate element boundaries through a statistical approach, allowing the identification of significantly co-occurring left and right terminal signal pairs. We benchmarked HELIANO against the manually curated HLEs database of the *Fusarium oxysporum* genome, and we then used it to perform an in-depth prediction of HLE in three large genomes. Finally, we applied our new tool to scan the genomes of 404 eukaryotic species spanning the whole Tree of Life. We further annotated all predicted HLEs for their additional domains, built a new, largely extended phylogenetic tree of HLEs and proposed new perspectives on HLE classification. HELIANO is more accurate than previous HLE-annotation tools, is well-suited for large-scale, systematic analysis of HLEs in eukaryotes, and will thus be a valuable tool to further our understanding of HLE evolution and impact.

## Material and Methods

### Curation of *Helitron*-like elements from Repbase

Before detecting HLEs in genomes, we reasoned that having a global view of their structural features based on previously characterized elements would be helpful. We thus used four parameters to obtain a detailed description of the structural features of HLEs available in Repbase (27): the whole length of HLEs, the distance between Rep and Hel domains (d-RepHel), the distance between LTS and Rep domain (d-LTSRep), and the distance between Hel and RTS domain (d-RTSHel). We found that HLEs varied greatly in size from 53 nt to 39,893 nt, but about 75% of autonomous HLEs were shorter than 12,338 nt with an average length of 9,666 nt, while 75% of non-autonomous HLEs were shorter than 2,619 nt with an average length of 2,049 nt (Supplementary Figure 1A). The d-RepHel was shorter than 973 nt for about 75% of autonomous HLEs, with a maximum value of 2,439 nt (Supplementary Figure 1B). Moreover, we observed that the d-LTSRep value was shorter than 5,275 nt for 75% of autonomous HLEs with a maximum value of 27,501 nt, and the d-RTSHel value was shorter than 3,301 nt for 75% of autonomous HLEs with a maximum value of 16,811 nt (Supplementary Figure 1C, D). Based on the distribution of these HLE structural features, we set the default value of ‘dm’ in HELIANO program as 2,500, which is corresponding to the parameter d-RepHel; the default value of ‘w’ in HELIANO program as 10,000, which is corresponding to parameters d-LTSRep and d-RTSHel.

HLE variants differ in their terminal structure and coding potential (12). On one hand, these differences could be used to classify different variants. On the other hand, this requires different strategies to identify such variants. Although Repbase is a well-curated TE reference database where hundreds of autonomous HLEs have been collected, most of these collected HLEs have not been further classified into variants (27). To improve the annotation of HLEs in Repbase, we initially collected Rep and helicase protein sequences from previous studies where HLE variants had been classified (13, 22, 28). Because these sequences represented a small subset of HLEs, we expanded this dataset by searching homologous sequences in Repbase (27) using NCBI blastp with default parameters (v2.13.0+) (29). Including the query sequences from previous studies, we finally collected 239 helicase sequences: 167 for *Helitrons*, 72 for *Helentrons*, and 228 Rep endonuclease sequences: 155 for *Helitrons*, 73 for *Helentrons*. This expanded dataset represented a large diversity of species, including 13 fungi species, 20 land plants, 39 animals, two algae, and three Oomycota. Each homologous sequence found in Repbase was then classified into a specific variant based on the highest blastp score obtained. The classification of collected HLE sequences was further checked and curated through a phylogenetic analysis. We computed multiple alignments using mafft (v7.475) with the parameter ’--auto’ (30), inferred phylogenetic tree using FastTree (v2.1.11 with default parameters) (31), and removed ambiguous leaves for which the phylogenetic position was inconsistent with the classification determined by blastp results. We provided the classification information in Supplementary Table S1. We observed in the phylogenetic trees that the transposase of *Helitrons* and *Helentrons* were distinctly separated, suggesting a reasonable classification (Figure 1B, Supplementary Figures 2-3). Finally, we used this dataset to build the HMM model of HLE transposase used in HELIANO.

### Training HMM models for Rep and Helicase domains

We computed multiple alignments for Rep and Helicase sequences for each of the three HLE variants using mafft with the ’--auto’ parameter. We then ran hmmbuild (v3.3) with the default parameter on each aligned file to obtain the four HMM models used in HELIANO (Supplementary material) (32).

### Algorithm of HELIANO

The HELIANO program follows a simple strategy: the first step is to search autonomous HLEs based on the transposase amino-acid sequence motifs, and the second step is to identify their non-autonomous derivatives. We divided the pipeline into three main parts: transposase detection, LTS-RTS pair identification, and filtration (Figure 2). HELIANO relies on the prediction of ORFs in the genome sequence query and applies our pre-built HMM models to search for HLE joint Rep and Hel domains to find the transposase sequences (Figure 2A). HELIANO then scans the flanking region of transposases to identify significantly co-occurring LTS and RTS pairs (Figure 2B). Finally, HELIANO refines the candidates by checking the alignments of each subfamily’s 50 nucleotides (nt) flanking sequence containing identical LTS and RTS pairs. (Figure 2C). Here, as previous work suggested, we define families based on their RTS sequences and subfamilies based on their LTS sequences (12). The source code of HELIANO is available from zenodo (https://zenodo.org/records/10625240) and github (https://github.com/Zhenlisme/heliano/).

**Figure 2.**
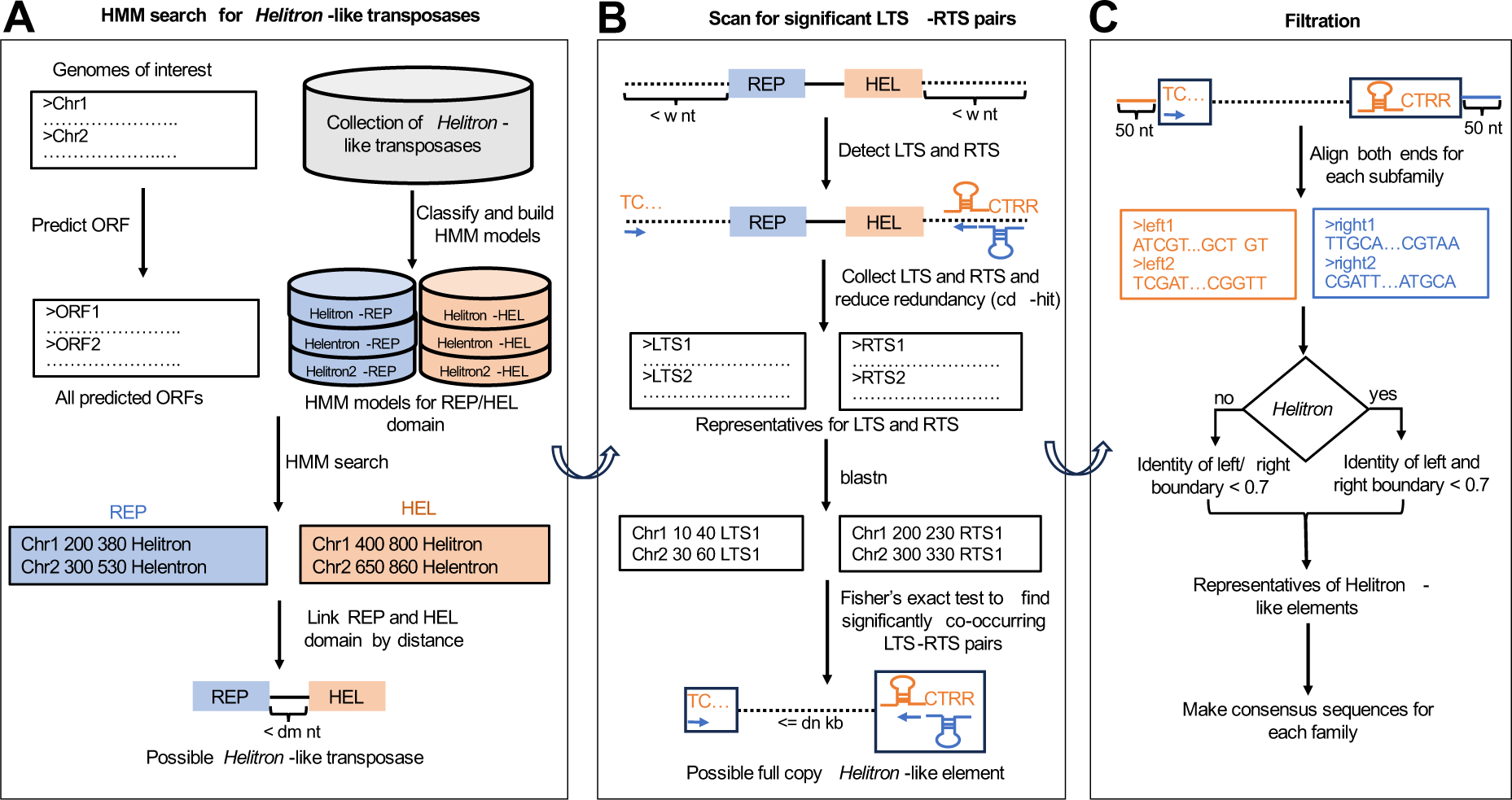
Workflow of HELIANO. (A) HMM searches for transposases of HLEs*; dm* denotes the distance between the Rep and Helicase domains. (B) Scan for significantly co-occurring LTS-RTS pairs; *w* indicates the length of the RepHel domain flanking sequences; *dn* denotes the distance between LTS and RTS. (C) Filtration to get representative insertions and make their consensus sequences.

### Search for potential HLE transposases in genomes

For a given genome, HELIANO uses the program getorf (EMBOSS:6.6.0.0) to predict open reading frames (ORF) with the parameter ’-minsize 100’ (33). Then HELIANO uses each trained HMM model to predict possible Rep endonuclease and helicase ORFs in the genome using hmmsearch (v3.3) with the parameter ’—domtblout --noali -E 1e-3’. To avoid false positives, HELIANO filters out hits with ’c_Evalue’ or ’i_Evalue’ lower than 1e-5. The classification of every hit as *Helitron* or *Helentron* is further determined by selecting the variant with the highest ’full-sequence score’. Because all HLE transposases contain both the Rep endonuclease domain and the Helicase domain, and because the Rep endonuclease domain is always upstream of the Helicase domain, we can deduce the genomic regions of *Helitron*-like transposase. HELIANO uses bedtools window (v2.30.0) to find the genomic coordinates for the Rep-Helicase region (34). The potential *Helitron*-like transposase is then classified based on the classification of Rep and Helicase domains (Figure 2A).

### Detection of LTS and RTS of HLEs

HELIANO then scans two windows at both ends of *Helitron*-like transposases up to a given distance (default is 10,000 nt). The left terminal sequences (LTS) are searched in the left or upstream window, and the downstream right terminal sequences (RTS) are searched in the right or downstream extended window. For the *Helitron* variant, HELIANO applies the LCV model developed by HelitronScanner to detect the LTS, which starts with the dinucleotide TC (8). The RTS is expected to form a stem-loop structure containing a ’CTRR’ motif, and this structure is searched using rnabob (v2.2.1). For *Helentron*, HELIANO uses tirvish of the GenomeTools package (v1.6.1) (35) to detect TIRs whose left and right pairs will be taken as LTS and RTS, respectively. We set the size of TIRs to be longer than 11 nt and shorter than 18 nt according to previous studies (Figure 2B) (15, 16).

### Identification of autonomous and non-autonomous HLE*s*

For all transposase regions, HELIANO collects sequences from all detected LTS and RTS sequences with an extension of 30 nt. These sequences are clustered to obtain unique sets of LTS and RTS using cd-hit (v4.8.1) (36). Next, HELIANO searches all homologous sequences using unique LTS and RTS sequences as queries against the genome using NCBI blastn (v 2.13.0+) (bitscore >= 32 by default). Finally, HELIANO retrieves all LTS-RTS pairs whose LTS should be upstream of its RTS using bedtools (v2.30.0) (34). By default, HELIANO searches LTS-RTS pairs whose LTS and RTS originate from the same transposase. To identify the best LTS-RTS pair, we test whether the sets of LTS and RTS sequences of each pair colocalize in the whole genome, i.e. are located less than *dn* bp apart from each other. HELIANO takes advantage of the Fisher’s exact test wrapped in the program bedtools to find such pairs. LTS and RTS sequences that are significantly co-occurring in a given genome would be taken as potential terminal sequences of HLEs, including autonomous and non-autonomous copies. We further classify these pairs into families based on their RTS sequences and subfamilies based on their LTS sequences. For example, two candidates could be classified into the same family if their RTS sequences share at least 90% identity. The candidates from the same family could be further classified into the same subfamily if their LTS sequences share at least 90% identity (Figure 2B).

### Selection of the representative candidates from all possibilities

Inner LTS-RTS pairs existing within the intervals defined by the selected LTS-RTS pairs can also pass Fisher’s exact test introduced in the last step. To examine such cases, for each subfamily, HELIANO samples up to 20 sequences, including 50 nt of flanking residues and performs a multiple alignment using mafft. We reasoned that flanking residues are conserved if they belong to the transposon while flanking residues of the real LTS-RTS are not expected to be conserved. HELIANO evaluates the average identity of aligned sequences using the R package seqinr (v4.2.30) (37). Ultimately, subfamilies with less than 70% identical flanking regions are selected as representative *Helitron*-like element candidates, while the remaining ones constitute alternative candidates (Figure 2C).

### Benchmarking

We needed a reliable database as ’ground truth’ for benchmarking HELIANO and the other tools for HEL annotation. We identified the study of Chellapan and collaborators as suitable for this benchmarking because they manually curated HLEs in ten *F. oxysporum* genomes (15). As a result, they characterized five families of *Helentron* variants and 26 consensus sequences that can be found in Repbase. We selected the genome of the Fo5176 strain (GCA_030345115.1) for benchmarking because it represents the most contiguous *F. oxysporum* genome with 4.5 Mbp for N50, 7 Mbp for L50, and 70.1 Mbp for genome size. To ensure that all complete insertions were fully recovered, we collected all LTSs and RTSs of *Helentrons* described in that study: 25 unique LTSs and 24 unique RTSs (15). Next, we used blastn using an e-value cutoff of 1e-2 to find all their homologous sequences in the Fo5176 genome. Then, we recovered full insertions by pairing all LTSs and RTSs using the window function of bedtools. After manual curation, we finally recovered 253 full insertions (Supplementary Table S2), used as a ’ground truth’ in the following benchmarking process. HELIANO was run with the parameter ‘-w 15000 -is2 0 -p 1e-5 -n 20’; HelitronScanner was run with the default parameters; EAHelitron was run with the parameter ‘-r 4 -p 20 -u 20000’; RepeatModeler2 (v2.0.5) was run with the default parameter (8, 26, 38). As RepeatModeler2 only outputs consensus sequences, we then recovered their corresponding full copy insertions (>= 80% length of consensus) with the blastn program. We executed each program using a computer operated under Ubuntu GNU/Linux 22.04 LTS system with 20 threads and reported the real time of execution.

We then designed four benchmarking matrices for each program to evaluate their performance, including precision, sensitivity, FDR, and F1 score, computed using standard formulae (39). True positive (TP) was defined as the number of predicted insertions with more than the cutoff overlap in length with real insertion. The remaining predicted insertions were defined as false positives (FP). False negative (FN) was defined as the number of real insertions with less than the cutoff overlap in length with any predicted insertions. Eight cutoffs were further tested to calculate FP, FN, and TP: 65%, 70%, 75%, 80%, 85%, 90%, 95%, and 100%.

### Comparison between HELIANO prediction and Repbase dataset for *X. tropicalis, X. laevis* and *O. sativa*

We downloaded the *Xenopus tropicalis* (GCF_000004195.4) and *Xenopus laevis* (GCF_017654675.1) genomes from NCBI, the *Oryza sativa* genome (version 7.0) from the RGAP website (http://rice.uga.edu/). For *X. tropicalis* and *X. laevis* genome, we ran HELIANO with the parameter ‘-s 30 -is1 0 -is2 0 -sim_tir 90 -n 20 -p 1e-5’. For *O. sativa* genome, we ran HELIANO with the parameter ‘-s 30 -is1 0 -is2 0 -n 20 -p 1e-5’. To assess the precision of HLE boundaries annotated by HELIANA, we manually examined each HLE subfamily’s insertions by aligning the predicted insertion sequence with its genomic loci using *dialign2* (40). For *O. sativa* and *X. tropicalis*, we searched homologous sequences using the corresponding Repbase HLE consensus as queries against their genomes using NCBI blastn. We identified complete copies from the hits that shared at least 80% identity and 80% coverage to the query. These full-copy datasets were named Rbfull-XT for *X. tropicalis* and Rbfull-OS for *O. sativa* (Supplementary Table S5 and S6). We then used bedtools intersect to compare HELIANO and Rbfull insertions. An insertion was considered present in Rbfull and HELIANO if the Rbfull insertion was covered by at least 80% of its length.

### HELIANO annotation on selected genomes

We established a selection of eukaryotic genomes as follows: 1) we downloaded the taxonomic information from 2,302 species whose genome assembly level reached the chromosomal level from the NCBI assembly database. 2) we then randomly selected the species by ensuring every order has at most two species, which will not represent the same family or genus. We removed the species *Trichomonas stableri* from the list because we could not find its genome assembly in NCBI. As a result, we finally identified 404 well-assembled genomes (Supplementary Table S3). We then ran HELIANO with default parameters for each of these sampled genomes. Four thousand four hundred ninety-one bacterial genomes downloaded from NCBI were also tested as negative controls (Supplementary Table S4).

### Detection of additional protein domains in HLEs from sampled genomes

For each detected autonomous HLE, we used the program getorf (EMBOSS:6.6.0.0) to predict their ORFs with the parameter ‘-minsize 100’. Then we used hmmsearch to identify additional domains in HLEs with the parameter ‘-E 1e-3’ from Pfam downloaded from the InterPro website https://ftp.ebi.ac.uk/pub/databases/Pfam/current_release/Pfam-A.hmm.dat.gz (41). We reasoned that a domain captured by a family of HLE should be found in most copies belonging to the same HLE family. Moreover, we expected that random TE insertions could contribute to domains in HLEs, and we needed to exclude such cases from our analysis. To do so, the first filter we used was to remove domains found in less than five or 50% of HLE copies. Since many domains remained scattered along the whole length of HLEs, we empirically determined that filtering out the domains that were more than 4,000 bp away from the RepHel domain and removing the domains that occurred upstream and downstream of RepHel proved effective. Because of the lack of specific annotation for HLE transposases in Pfam, their Rep and Helicase domains have been annotated as different Pfam families. The Rep domain was annotated as families of Helitron_like_N (N-terminal of HLE transposase) and RepSA (replication initiator protein). The Helicase domain was annotated as families of PIF1/Pif1_dom_2B (Pif1-like Helicase). We thus ignored RepSA domains from the result and merged the name of Pif1_dom_2B with PIF1.

### Construction of phylogenetic trees of HLEs from genomes of sampled species

For each species whose HLEs could be detected by HELIANO, we used the program getorf (EMBOSS:6.6.0.0) to predict ORFs of all its HLEs with the parameter ‘-minsize 400 -find 1’. Then, all Rep and Helicase domains were predicted via hmmsearch (v3.3) with the parameter ‘-E 1e-3’ based on the same hmm model used in HELIANO. Predicted Rep and Helicase amino acid sequences of each HLE were extracted and concatenated into single sequences (RepHel). For each species, we then ran the program cd-hit with the parameter ‘-c 0.7’ to get the representatives of RepHel sequences. An outgroup sequence was made by concatenating the geminivirus rep catalytic protein (NCBI accession number: WP_015060107.1) and helicase protein of *Myroides phaeus* (NCBI accession number: WP_090404604.1). All representatives of RepHel sequences and the outgroup sequence were aligned using mafft (v7.475). Finally, FastTree (v2.1.11) was applied to reconstruct a phylogenetic tree with default parameter (31).

## Results

### HELIANO benchmarking and comparison with other tools

We used the published *Helentron* dataset of *F. oxysporum* as a ’ground truth’ for benchmarking (15). We selected it because it is the only accessible manually curated dataset, as far as we know. We recovered 253 full *Helentron* insertions, which were taken as genuine in the following benchmarking process (15). To estimate the performance of HELIANO, we calculated its precision, sensitivity, FDR, and F1 under different overlap cutoffs. We evaluated the performance of HelitronScanner, EAHelitron and RepeatModeler2 using the same method (Figure 3).

**Figure 3.**
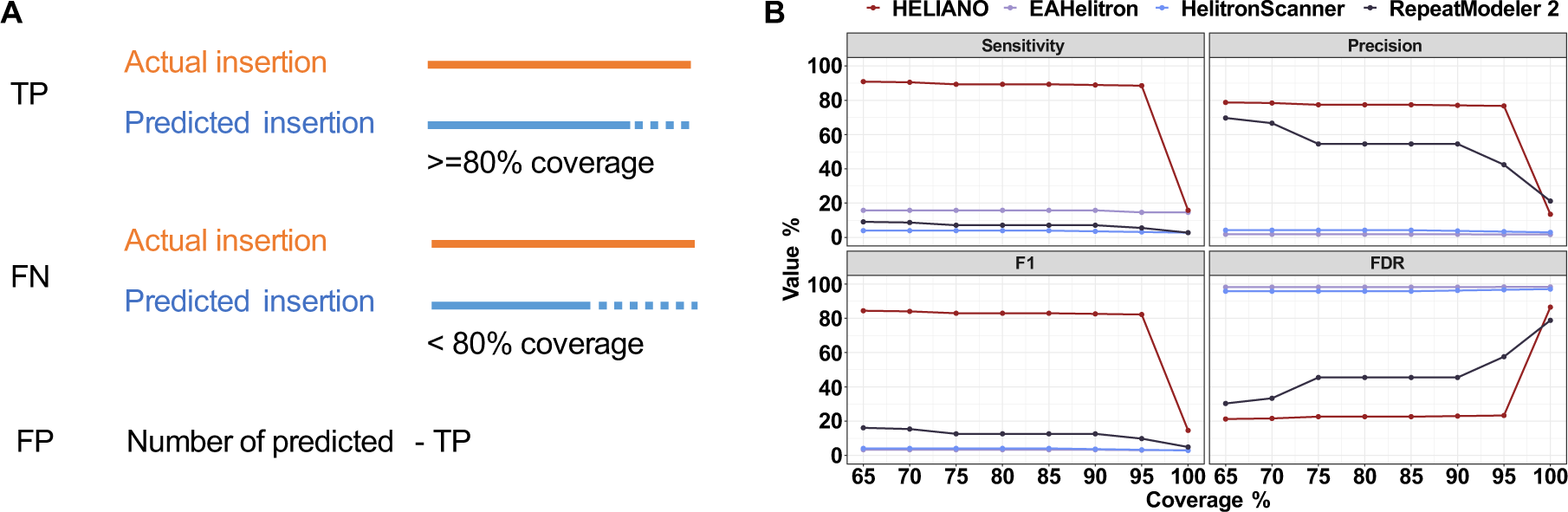
Benchmarking analysis of HELIANO. (A) Schematic representation of benchmarking metrics. TP: True positive; FN: False negative; FP: False positive. (B) Comparison of benchmarking metrics of all tested software. F1 is the score computed as the harmonic mean between sensitivity and recall. FDR: False Discovery Rate.

HELIANO had the highest sensitivity and F1 score among all test software (Figure 3B). HELIANO could detect 224 of the 253 genuine insertions (88.53%) with a coverage cutoff of 95%. EAHelitron found 40 insertions (15.81%) at the same cutoff level. Similarly, RepeatModeler2 identified 14 insertions (5.53%), and HelitronScanner only had eight (3.16%, Supplementary Table S2). Overall, HELIANO had the lowest FDR and the highest precision. HELIANO predicted 68 complete insertions absent in the genuine insertion dataset, with an FDR of 23.29% at a 95% cutoff level (Supplementary Table S2). We found that most of these predictions were characterized by clear terminal signals, indicating they might be new families of HLEs that have not been discovered yet (Supplementary Table S2). The EAHelitron software annotation had the highest FDR value, with 98.30% insertions absent in the genuine dataset. The software RepeatModeler2 had the second highest precision (69.70%), close to HELIANO at a 65% cutoff level. But when the cutoff was increased, RepeatModeler2 precision reduced while HELIANO kept a better precision level (76.71% ∼ 78.77%).

Regarding the execution time, RepeatModeler2 was the slowest software, with about two and a half hours, likely because it annotates all TEs. HelitronScanner took 35 minutes and EAHelitron 92 seconds. HELIANO ran the fastest with 70 seconds.

### HELIANO uncovers overlooked HLEs in *Xenopus* frog genomes

As a first test case, we ran HELIANO on two frog genomes to further annotate their HLEs. The pipid frogs *X. tropicalis* and *X. laevis* are two important vertebrate model species with high-quality chromosomal scale genome assemblies in which TEs have been annotated (42, 43). Moreover, these frog genome sequences are large and complex: 1.4 Gbp for *X. tropicalis* and 2.7 Gbp for the allotetraploid *X. laevis*, and their TE landscape is characterized by a majority of class II TE (42, 43). Only three non-autonomous *Helentron* consensus sequences have been reported for *X. tropicalis,* and none has been described for *X. laevis* in Repbase or previous studies (42). We annotated HLEs in these genomes using HELIANO in five minutes and 59 seconds for *X. tropicalis* and 16 minutes and 32 seconds for *X. laevis*.

Based on the 80-80 rule, we could map back the three Repbase non-autonomous *Helentron* sequences in the *X. tropicalis* genome and identified 638 insertions (Rbfull-XT dataset, Supplementary Table S6). Using HELIANO, we annotated 82 *Helentron* insertions and no *Helitron* in the *X. tropicalis* genome. These *X. tropicalis* insertions included three autonomous and 79 non-autonomous *Helentrons*, further classified into three families based on the RTSs homology (Table 1, Supplementary Table S6). About 97% (72 out of 74) of HELIANO-specific predictions belonged to the same family, HelenXT233. We found that only eight non-autonomous HLE insertions annotated by HELIANO were also in the Rbfull-XT dataset. These insertions were annotated as the HelenXT233 family in HELIANO prediction and the Helitron-N2_XT family in the Rbfull-XT dataset. However, we did not find the HELIANO-prediction counterparts for the other two Rbfull-XT families, Helitron-N1_XT (564 insertions) and Helitron-N1A_XT (59 insertions), which together made about 99% of the remaining Rbfull-XT-specific insertions. As expected this difference stemmed from HELIANO’s inability to detect families whose autonomous HLEs are absent from the genome.

**Table 1.** HELIANO-predicted and Repbase full copies of HLEs in *Xenopus* genomes.

| Species | Family | Subfamily | Auto | Nonauto | Variant | Source |
| --- | --- | --- | --- | --- | --- | --- |
| *X. tropicalis* | HelenXT233 | 2 | 1 | 79 | *Helentron* | HELIANO |
| *X. tropicalis* | HelenXT102 | 1 | 1 | 0 | *Helentron* | HELIANO |
| *X. tropicalis* | HelenXT365 | 1 | 1 | 0 | *Helentron* | HELIANO |
| *X. tropicalis* | Helitron-N1_XT | - | 0 | 564 | *Helentron* | Repbase |
| *X. tropicalis* | Helitron-N1A_XT | - | 0 | 59 | *Helentron* | Repbase |
| *X. tropicalis* | Helitron-N2_XT | - | 0 | 15 | *Helentron* | Repbase |
| *X. laevis* | HeliXL45 | 2 | 8 | 408 | *Helitron* | HELIANO |
| *X. laevis* | HeliXL2 | 2 | 3 | 1238 | *Helitron* | HELIANO |
| *X. laevis* | HelenXL108 | 1 | 1 | 545 | *Helentron* | HELIANO |
| *X. laevis* | HelenXL83 | 1 | 1 | 0 | *Helentron* | HELIANO |
| *X. laevis* | HelenXL460 | 1 | 2 | 7 | *Helentron* | HELIANO |

Although there were no HLE sequences ever reported In the *X. laevis* genome (27, 42), HELIANO annotated 2,213 full insertions, including 15 autonomous and 2,198 non-autonomous insertions, which can be classified into three *Helentron* and two *Helitron* families (Table 1, Supplementary Table S7).

We manually examined each HLE subfamily’s insertions by aligning the predicted insertion sequence with its genomic loci. We found that HELIANO annotated correctly their near full-length insertions of autonomous and non-autonomous *Helitrons* and *Helentrons*. Their boundaries could be confirmed by their T-T insertion sites for *Helentrons* and A-T for *Helitrons* and by the precise alignment at both terminal regions and discordant alignments in flanking regions (Figure 4C, D, E). We could identify each family’s terminal features, such as the TC motif in the LTS and stem-loop with CTRR suffix for *Helitrons* and the terminal inverted repeats and stem-loop structures in RTSs for *Helentrons*. (Figure 4C, D, E). But for some families, like HelenXT102 and HelenXT365, we failed to identify their insertion sites, and their boundaries were hard to find, which might represent degenerated *Helentron* insertions. Moreover, we found that the terminal sequences of the autonomous insertion HelenXT233 were almost identical to those of Helitron-N2_XT, described in Repbase for decades, while its autonomous origins had never been discovered. This further evidenced the robustness of HELIANO for identifying autonomous HLEs and their non-autonomous derivatives in the large and complex *Xenopus* genomes (Figure 4C, D).

**Figure 4.**
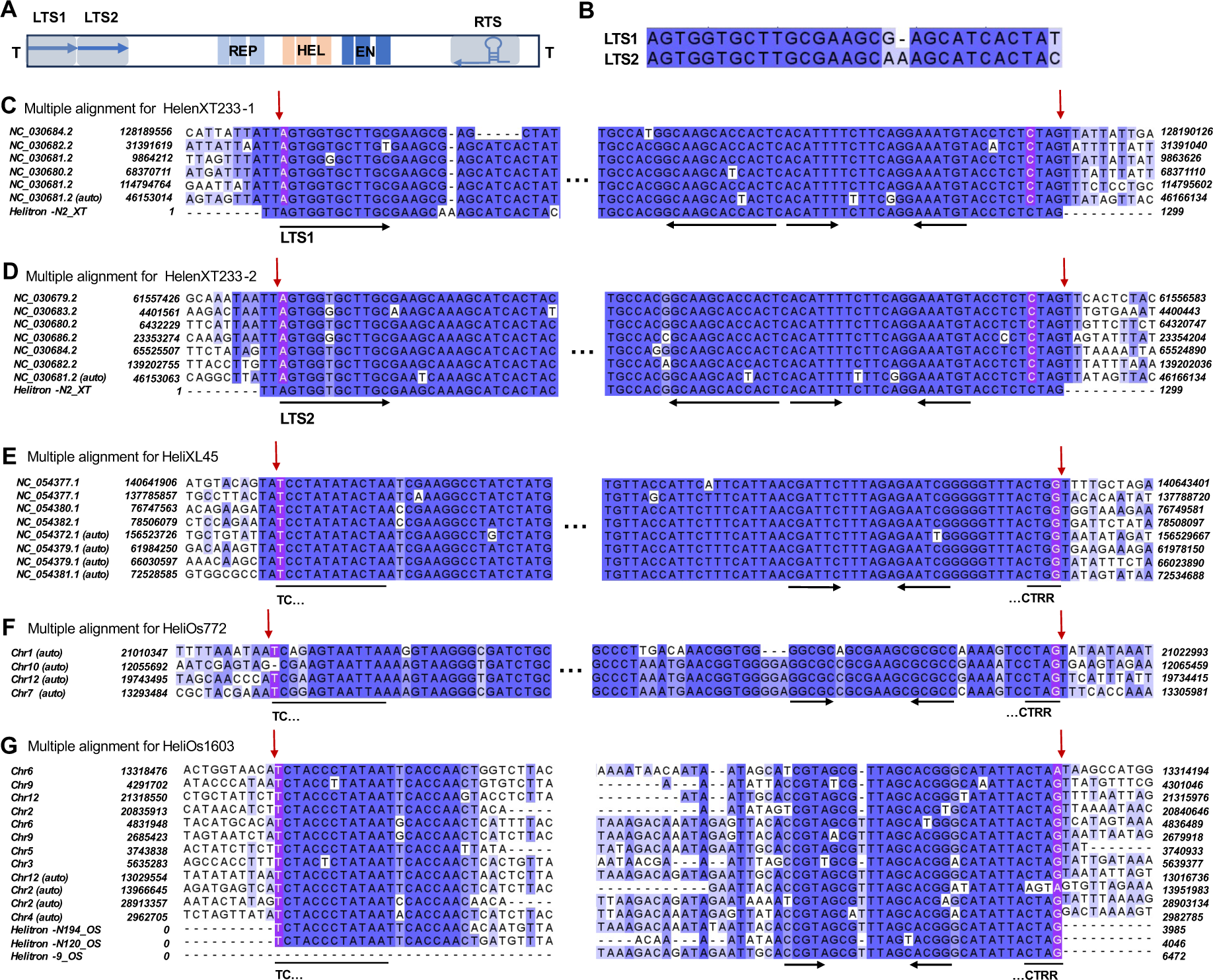
Multiple alignments of selected *Helitron* and *Helentron* insertions detected by HELIANO for *Xenopus* frog and *Oryza sativa* genomes. (A) Structure of the *X. tropicalis* autonomous HelenXT233. There are two alternative LTSs detected: LTS1 and LTS2. (B) Sequence alignment of LTS1 and LTS2 from the autonomous *X. tropicalis* HelenXT233. (C, D) Multiple alignments of insertions from HelenXT233 families (C) for HelenXT233-1 and (D) for HelenXT233-2. (E) A case for multiple alignments of *X. laevis Helitron* insertions. (F, G) Cases of multiple alignments of *Helitron* insertions in *O. sativa* genome. The autonomous insertions are labelled as ’auto’ in each multiple alignment, and others are non-autonomous counterparts. The nucleotide highlighted in purple shows the predicted starts and stops by HELIANO. The down arrows in red indicate the precise insertion sites based on manual curation, using as a rule that *Helentrons* are inserted between T and T nucleotides and *Helitrons* are inserted between A and T nucleotides. Note the precise correspondences between the HELIANO annotation and the manual curation for *Helentrons* in E-F-G and the differences for *Helitrons* in C and D. The horizontal black arrows indicate terminal inverted repeats and stem-loop structures.

It is well known that HLE LTSs are more diverse than their RTSs (8, 12). We made similar observations for HLEs in *Xenopus* genomes. For example, the families HelenXT233, HeliXL45 and HeliXL2 could be further classified into subfamilies based on their LTSs homology (Table 1). The autonomous HelenXT233 is characterized by two LTSs, LTS1 and LTS2, forming a direct repeat (Figure 4A). However, LTS1 and LTS2 are not identical, each hallmark a different subfamily of non-autonomous *Helentrons*. LTS1 characterizes helenXT233-1, and HelenXT233-2 is characterized by LTS2 (Figure 4B, C, D).

### HELIANO uncovers overlooked HLEs in *O. sativa*

As a second test case, we ran HELIANO on the *O. sativa* genome, one of the most important plant models (44, 45). HELIANO ran the task in 18 minutes and 26 seconds. There are 310 *Helitron* consensus sequences collected from *O. sativa* in Repbase, including 22 autonomous and 288 non-autonomous entries. From these, we could map back only 14 autonomous HLEs consensus autonomous sequences and 236 non-autonomous ones in the genome sequence based on the 80-80 rule (2). These consensus sequences contributed to 25 autonomous and 2,088 non-autonomous complete insertions in the rice genome (RbFull-OS dataset, Supplementary Table S5). We ran HELIANO on this rice genome and predicted 79 autonomous and 1769 non-autonomous *Helitron* insertions without evidence for *Helentron* (Supplementary Table S8).

We then asked how many insertions annotated by HELIANO could also be found in the Rbfull-OS dataset. HELIANO annotated 21 autonomous insertions out of 25 (84%) in RbFull-OS. We examined the four autonomous insertions that differed and found that they were annotated in the HELIANO dataset but did not match over their entire length with their RbFull-OS equivalent, indicating that HELIANO predicted different LTS and RTS. We asked if the 58 HELIANO-unique autonomous insertions were new predictions or drawbacks of homology searching methods used to compile the RbFull-OS dataset. We ran a phylogenetic analysis to compare these 58 HELIANO-unique predictions and all 14 RbFull-OS autonomous *Helitrons*. The result showed that while all RbFull-OS autonomous *Helitrons* had identical counterparts in HELIANO prediction, the converse was not true, i.e. HELIANO unveiled new insertions (Supplementary Figure 4). We confirmed this finding by clustering all these HLE insertion sequences at the 90% identity threshold and observing that 44 clusters were HELIANO-specific (Supplementary Table S9). For example, HELIANO successfully annotated the total insertions of family HeliOs772 absent in RbFull-OS (Supplementary Figure 4, Supplementary Table S8). Its actual insertions were confirmed by their canonical *Helitron* structures (started with TC dinucleotide and stopped with stem-loop structure with CTRR suffix) and insertion sites between A and T (Figure 4F).

For non-autonomous HLEs, HELIANO and RbFull-OS results differed significantly since 1,796 insertions were RbFull-OS-specific and 1,484 HELIANO-specific. The HELIANO-specific insertions could be explained by the fact that RbFull-OS included only a few non-autonomous complete copies for each family. At the same time, by design, HELIANO recovers non-autonomous insertions corresponding to autonomous ones based on shared LTS and RTS. For example, HELIANO predicted four autonomous and 60 full non-autonomous insertions for the family HeliOs1603 that had only one autonomous (Helitron-9_OS) and six non-autonomous counterparts in RbFull-OS (Figure 4G, Supplementary Figure 4). About 75% of HELIANO-predicted HeliOs1603 insertions were shorter than 7 kb, indicating that they were not likely to be false positives. The RbFull-OS-specific insertions could be attributed to the absence of their autonomous counterparts in the genome because HELIANO, by design, can not detect non-autonomous HLEs whose autonomous counterparts are missing.

We concluded that HELIANO could be especially useful for classifying and identifying *Helitrons* in complex and HLE-rich plant genomes such as rice.

### HELIANO revealed a broad distribution of *Helitrons* and *Helentrons* in the eukaryotic world

To further evaluate the applicability of HELIANO, we sampled 404 chromosome-level genome assemblies of eukaryotic species from the NCBI genome database. The tested genome size ranged from 7.30 Mbp to 40.05 Gbp, and their GC content varied from 16.59% to 78.37% (Supplementary Figure 5). The corresponding species list covered 27 phyla, 83 classes, and 281 orders, including fungi, animals, land plants, and algae (Supplementary Table S3). Thus, this dataset represents a wide range of genome complexity and species diversity. We included 4,491 bacterial genomes expected to lack HLEs and used them as a true negative dataset according to the current model on HLE evolution (Supplementary Table S4) (46, 47).

We did not detect any HLEs in the sampled bacterial species, while HLEs were widespread among eukaryote genomes (Figure 5A). In addition, we found that 139 species lack HLEs in their genomes. For example, HELIANO did not detect any HLE sequences among all 29 sampled bird genomes. Among 26 sampled mammalian genomes, we only found *Helitron* presence in the bat genome, as previously reported (19) (Figure 5A, B). Among the 404 tested eukaryote genomes, we identified 265 cases (66%) containing at least one HLE, encompassing 22 phyla and 61 classes (Supplementary Table S10, Figure 5A). Previous studies suggested a much narrower distribution of *Helentrons* than *Helitrons,* with a seeming absence in land plant genomes (12). However, in our large-scale scan, we found that both variants were prevalent all over the eukaryotic world. *Helitrons* were detected in 179 genomes from 20 phyla and 53 classes, *Helentrons* in 173 genomes from 19 phyla and 44 classes, and the unclassified HLEs in 19 genomes from eight phyla and 14 classes (Figure 5A, Supplementary Table S10). Furthermore, we identified a significant number of *Helentrons* in six (13 species) out of the eight (74 species) sampled land plant classes (Figure 5). We present the insertion of the HelenSM92 family in the *Sphagnum magellanicum* genome as an example of *Helentron* presence in land plants, where we observed *Helentron* features: short terminal inverted repeats and the ‘TT’ insertion sites (Supplementary Figure 5A). We also analyzed the protein domains of this HelenSM92 family using the Conserved Domain Database (CDD) search tool (48). Besides the RepHel domains, we found that a GIY-YIG domain was captured and transposed in this *Helentron* family (Supplementary Figure 6B-D).

**Figure 5.**
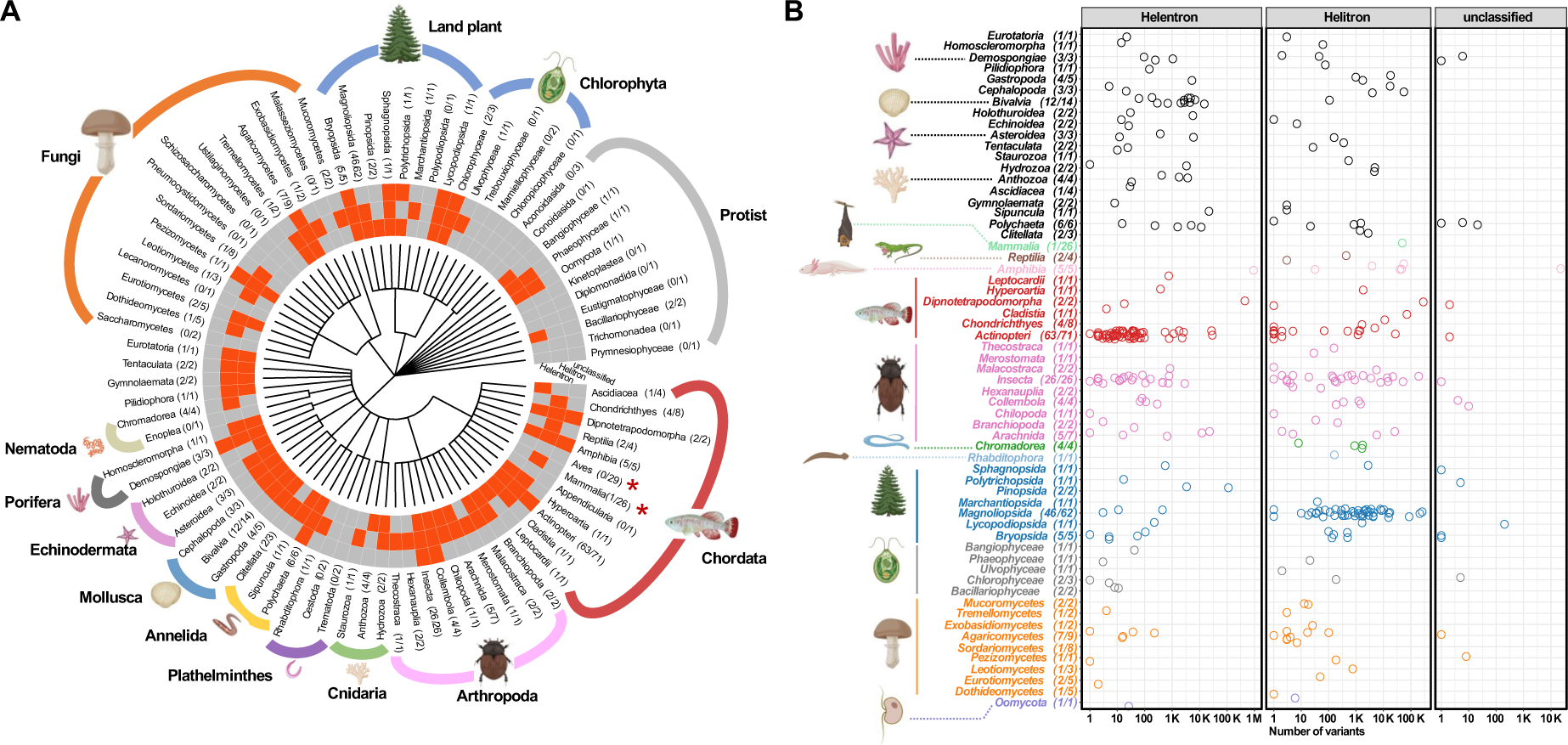
Distribution of HLEs among 404 eukaryote genomes. (A) A species tree obtained from NCBI indicated the phylogenetic relationship of sampled genomes. The fraction in each bracket represents the ratio of the number of species with HLEs to the number of all sampled species in a particular class. The heatmap indicates the presence (red) and absence (grey) of HLE variants in each sampled class. (B) The scatter plot shows the number of detected HLEs in each sampled class. Each point represents the number of corresponding HLE variants in a species. The fraction in each bracket represents the ratio of the number of species with HLEs to the number of all species in a particular class. The y-axis scale is log10 transformed.

We conclude that HLEs are widely distributed in eukaryote genomes and that the prevalence of *Helentrons* was underestimated in previous studies. Moreover, our results showed that HELIANO is a robust tool for annotating HLEs for complex and large genomes of diverse compositions.

### Capture of additional gene domains in HLEs

HLEs are well known for their ability to capture gene sequences (9, 49–51). Additional protein domains recurrently found in HLEs, such as the RPA, OTU and EN domains, were thought to originate from different ancient gene-capture events (12). To explore the potential gene-capturing events across HLEs, we annotated the protein domains for each detected autonomous HLE in each sampled species. We expected that all copies of the same HLE family would be characterized by a captured domain found at a conserved position if this domain was stably captured and transposed.

Besides the most recurrently detected *Helitron* helicase-like domain at N-terminus (Helitron_like_N) and PIF1 domains that we considered as being part of the HLE transposase, we found 20 additional domains that were stably included in HLEs, including the three previously described domains, RPA, OTU, and EN (annotated as Exo_endo_phos in Pfam) (12) (Supplementary Table S11). Overall, most of these 20 protein domains are known to enable the binding or modification of DNA or proteins. For example, the three amino acid peptide repeats (STPRs) and the B3 DNA binding domain (B3) function as transcription factors (52, 53). RPA and Ssb-like_OB are involved in DNA replication by binding to single-strand DNA (54, 55). The 2OG-Fe(II) oxygenase (2OG-Fell_Oxy_2) is reported to function as a DNA repair enzyme that removes methyl adducts and some larger alkylation lesions from endocyclic positions on purine and pyrimidine bases (56). The domain OTU, F-box associated domain (FBA_3), Ubiquitin carboxyl-terminal hydrolase (UCH), and C-terminal Ulp1 protease (Peptidase_C48) are involved in the regulation of protein degradation (57–61).

We further checked the position of these domains within HLE sequences. Globally, we found that all domains were enriched in certain regions, either downstream or upstream of RepHel, indicating their conserved position within HLEs (Figure 6). Some domains are likely to share the same open reading frame (ORF) with RepHel transposase, e.g., the domain DUF6570, Helicases from the Herpes viruses (Herpes_Helicase), N-terminal of large tegument protein of herpesviruses (Herpes_teg_N), and UvrD-like helicase C-terminal domain (UvrD_c_2) and provide evidence for molecular evolution events on the HLE transposase (Supplementary Figure 7A, 11, 12). However, the other 17 domains are encoded in different ORFs. Moreover, most domains were not shared between *Helentrons’* and *Helitrons’* variants. For example, the B3, protein phosphatase 2A regulatory B subunit (B56), FBA_3, Fn3-like domain from Purple Acid Phosphatase (fn3_PAP), Herpes_Helicase and STPRs were almost exclusively found in *Helitrons*. In contrast, 2OG-Fell_Oxy_2, DUF3106, DUF6570, EN, Herpes_teg_N, hemopoietic IFN-inducible nuclear protein (HIN), heat shock protein Hsp20 family (HSP20), Ssb-like_OB, and UCH were almost exclusively found in *Helentrons* (Figure 6). The remaining domains, e.g. RPA and OTU, were found in *Helentron* and *Helitron* variants (Figure 6). Interestingly, we found that the distribution features of RPA varied between *Helentron* and *Helitron*. The RPA domain was enriched downstream of RepHel in *Helitron* but upstream in *Helentron*, suggesting *Helitron* and *Helentron* might have independently captured the RPA gene (Figure 6). Previous studies did not detect the presence of OTU in *Helitrons* (12). However, we noticed this domain in both *Helitron* and *Helentron* upstream of RepHel (Figure 6; Supplementary Figure 7C-D, 8-9). Further phylogenetic analysis showed that the OTU domain of *Helitrons* was distinct from the OTU domains of *Helentrons* (Supplementary Figure 10). We conclude that gene capture events occurred repeatedly in HLEs and provided diverse molecular functions to these TEs.

**Figure 6.**
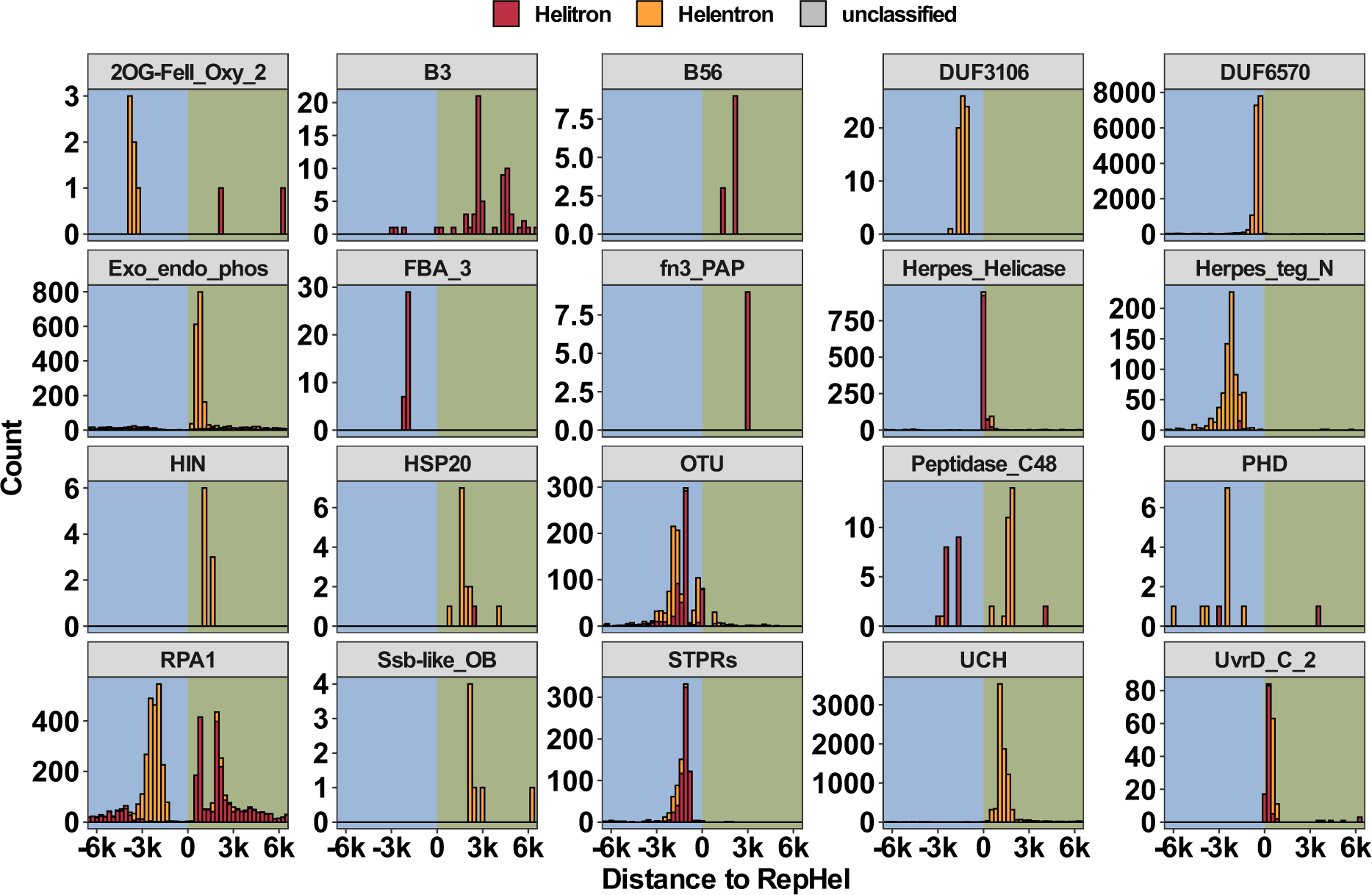
Distribution of the distance between RepHel and additional protein domains in HLEs. The zero value on the x-axis indicates the position of RepHel domains, negative values indicate the corresponding domains are upstream of RepHel, and positive value indicates their presence downstream of RepHel. The y-axis shows the count of HLEs.

### Phylogenetic distribution of HLEs in eukaryotic genomes

Previous studies suggested that HLEs could be classified into three different variants, *Helentron*, *Helitron* and *Helitron2,* based on the difference in their coding potential and structural features (12, 13). However, these analyses relied on a relatively small dataset (12, 13). Since we obtained a large number of HLEs from a wide diversity of genomes across the Tree of Life with HELIANO, we had the opportunity to study HLE diversity from a broader perspective. We reexamined this classification and further asked if we found additional variants of HLEs and what were their phylogenetic relationship. Our results showed that HELIANO accurately classified *Helitrons* and *Helentrons* following the phylogenetic classification. The accuracy of the HELIANO classification was estimated to be 99.17% (Figure 7A). We discovered divergent subgroups within the two main clades that classified all HLEs into *Helitrons* and *Helentrons* clades. The *Helentron* clade could be further classified into four subgroups (a, b, c, and d) and the *Helitron* clade could be further classified into five subgroups (e, f, g, h, and i) (Figure 7A). In addition to the nine subgroups, we found a few HLEs in at least four additional clusters. Known *Helitron* sequences from Repbase were found in all five *Helitron* subgroups. The previously identified variant *Helitron2* sequences from the *F. oxysporum* genome were found within *Helentron* subgroup a, indicating that *Helitron2* corresponds to a monophyletic subgroup of *Helentron* (Supplementary Figure 13, Figure 7A). Most Repbase *Helentron* sequences were found in subgroup b. Besides, Repbase HLEs were absent from subgroup d, indicating that HELIANO has identified a much broader diversity of HLEs than contained in the current Repbase collection. The subgroups c and d together comprised a unique *Helentron* clade consisting of a massive diversity of *Helentrons,* which mostly came from two species with giant genomes: the lungfish *Neoceratodus forsteri* (34.6 Gbp) and the newt *Ambystoma mexicanum* (28.2 Gbp). We found that about five HLE subgroups (the subgroups c, d, g, h, and i) are specific to their host types. For example, more than 90% of *Helitrons* in subgroups g, h, and i are hosted in diverse land plant species (Supplementary Figure 14). Conversely, we also observed a great variety of host types in some subgroups, highlighting the invasive nature of some HLEs. For example, at least four host types (Fishes, Mollusca, Cnidaria and land plants) could be found in subgroup b and five host types in subgroup f (Fishes, Arthropoda, Cnidaria and Amphibians and Mammals).

**Figure 7.**
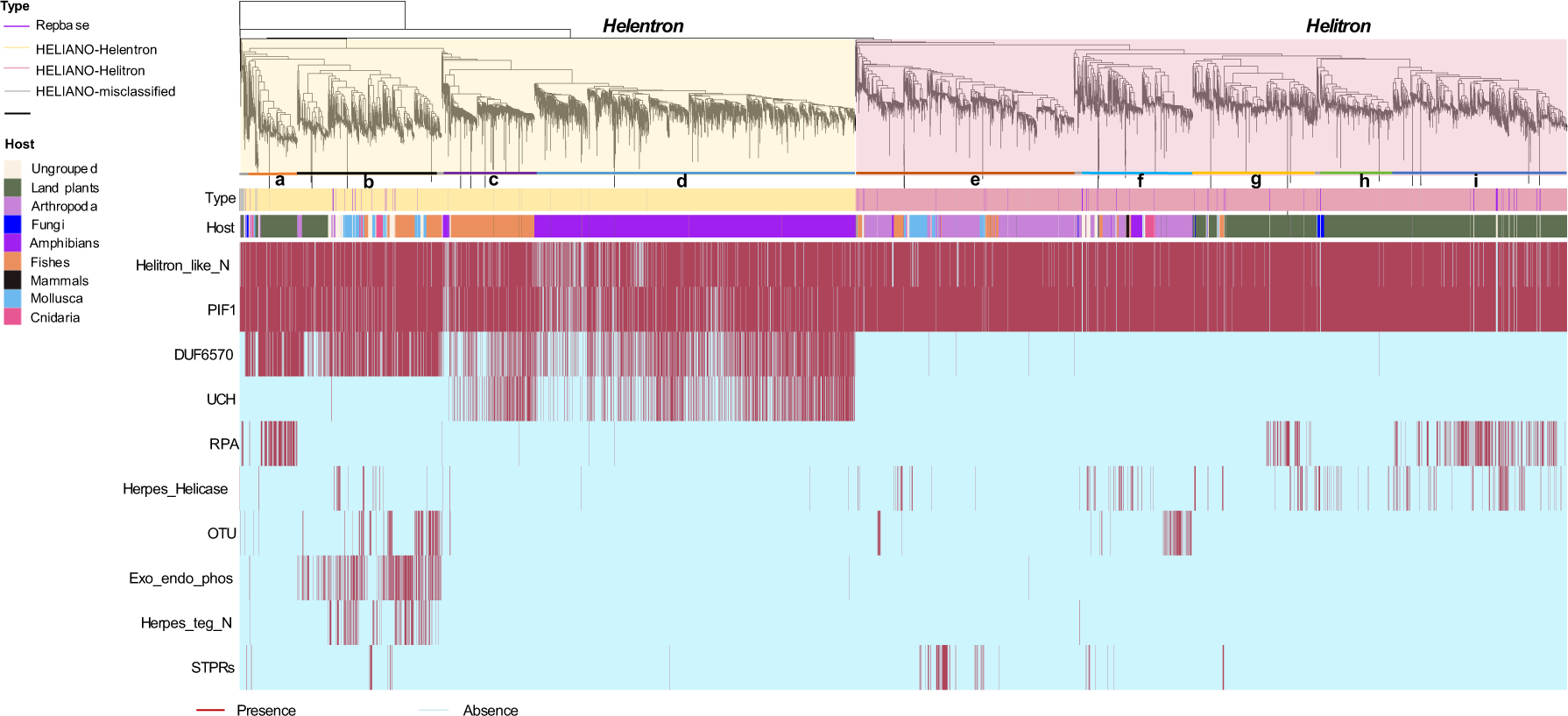
Distribution of HLEs and their captured domains in eukaryote genomes. (A) Maximum likelihood estimation tree of HLE transposases from sampled species (LogLk = -8062114.874). The *Helentron* (light yellow block) and *Helitron* (light red block) groups were further classified into subgroups: a-d for *Helentron* and e-i for *Helitron*. Unclassified HLEs are in grey. The “Type” heatmap below the tree indicates the classification and source of HLEs. HLEs from Repbase are marked in purple, in grey represent HELIANO misclassified HLEs, in light yellow represents HLEs predicted as *Helentron* by HELIANO, in light red represents HLEs predicted as *Helitron* by HELIANO. The heatmap entitled as Host represents the species origin of HLEs. (B) The heatmap shows the presence or absence of additional domains in each corresponding HLE. Red indicates the presence of the domain, and light blue indicates its absence.

We then asked if any additional domains within HLEs could be used as signals to classify them. We selected the top ten most frequently detected domains to analyze their distribution across the HLE phylogenetic tree. The RepHel domains represented by Helitron_like_N and PIF1 were included as positive controls (Figure 7 B). Within our expectations, RepHel were prevalent in all HLE clades. Globally, we found the domain DUF6570 specifically in the *Helentron* clade, suggesting it could be used as a marker to distinguish *Helitrons* from *Helentrons*.

Moreover, many other domains were found to be limited within specific subgroups. For example, the UCH domain was enriched in subgroups c and d in the *Helentron* clade, supporting their common origins (Figure 7B). The Exo_endo_phos and Herpes_teg_N domains were enriched in subgroup b. The RPA domain in *Helentron* was limited to subgroup a. All these examples suggested that gene domains in HLEs could be potential signals to understand HLE evolution.

## Discussion

Accurate TE annotation from genomic sequences is essential to genome annotation, especially in large eukaryote genomes (6, 38). Moreover, the accelerated pace of complete genome sequencing requires scalable methods to perform comparative analysis and to shed light on TE biology and evolution. Among DNA TE, HLEs stand out as relatively large mobile elements, ∼ 10 kbp, characterized by their ability to incorporate host gene DNA, but their annotation remains challenging (9, 23, 62).

Our HELIANO software provides a comprehensive solution to address HLE annotation in complete genomes, enabling large-scale comparative analysis. Due to the lack of species with a completely and perfectly annotated genome for HLEs, assessing the validity of HLEs annotation remains a complex task. In this study, we presented various analyses to support the relevance of HELIANO output. Using a manually curated set of HLE on the *Fusarium* genome, we show that HELIANO outperforms HelitronScanner, EAHelitron and RepeatModeler2. On this benchmark, our HELIANO software obtained the best precision, sensitivity, FDR and F1 metrics for all coverage values and was the fastest. While these results validated our algorithmic choices to develop HELIANO, they showed that full-length HLE annotation with precise boundaries at the base level is still challenging to obtain in some genomic loci.

On selected cases drawn from the analysis of two high-quality complex frog genomes, we showed the potential of HELIANO to predict *Helitrons* and *Helentrons* that were undetected so far. We found a strikingly different landscape of HLEs with autonomous and non-autonomous insertions in the diploid *X. tropicalis* and tetraploid *X. laevis* genomes that diverged 45-50 MYA (42, 43). Like HLEs annotated using *Fusarium* genomes, we could reproduce HELIANO performance on detailed annotation at the nucleotide-level resolution of TE boundaries, even though this depended on the genomic environment. To continue benchmarking, we targeted the *O. sativa* genome, a complex and HLE-rich plant genome (11). We identified 25 autonomous and 2,088 non-autonomous full insertions of HLEs in the rice genome based on existing annotations. We ran HELIANO and predicted 82 autonomous and 1,766 non-autonomous insertions. These annotation results gave the same picture of an HLE-rich genome dominated by non-autonomous transposons. HELIANO not only spotted all the autonomous *Helitrons* listed in Repbase but also uncovered many others. As expected, HELIANO predictions on non-autonomous HLEs were limited to families for which both autonomous and non-autonomous transposons were identified. Thus, *Helitron* annotation on the rice genome is a target for further methodological improvements, especially to detect non-autonomous HLEs for which no cognate autonomous elements exist.

Using a large set of eukaryote genomes, we found that HELIANO could quickly produce a range of annotations, from a lack of full-length HLE to predictions of thousands of full-length and non-autonomous copies. We did not predict any HLE using HELIANO on 4,491 bacterial genomes, in accordance with the current model on the evolution of HLEs (46). Similarly, we did not detect HLEs in 139 of the 404 screened eukaryote genomes, a finding underscoring that false positives are not a central issue. We found that both *Helentrons* and *Helitrons* were much more widespread across eukaryotes than expected. Among 27 sampled phyla, we identified *Helitrons* in 20 phylum genomes and *Helentrons* in 19 phylum genomes. Moreover, previous studies suggested that land plants lacked *Helentrons* (12). Yet, we verified *Helentron’s* presence in many land plant genomes (Figure 5A, 6A, Supplementary Figure 6, 14).

We also explored additional gene domains within the predicted HLEs. Besides the previously described domains (OTU, Exo_endo_phos, and RPA), we detected 17 more gene fragments incorporated into the HLE coding regions. These domains have various biochemical functions, such as transcription factors, protein degradation, etc. However, further work is required to investigate whether they are used in HLE transposition or involved in the host’s gene regulatory network. By checking the relative location of these domains to RepHel, we found that many domains had different distribution patterns between *Helitrons* and *Helentrons*, suggesting this information could be used as phylogenetic signals for their classification.

Further phylogenetic analysis about RepHel domains from all detected HLEs allowed us to re-examine the current phylogenetic classification of HLEs. Our results supported the existence of the two clades, *Helentron* and *Helitron* (Figure 6A). Besides, we identified four distinct subgroups in *Helentrons* (subgroup a-d) and five distinct subgroups in *Helitrons* (subgroup e-i). One previously described variant, *Helitron2,* was found within subgroup a, suggesting that *Helitron2* are only one of the four *Helentron* subgroups. Furthermore, we found that many subgroups were dominated by certain host types (Figure 6A), suggesting that the HLEs had the ability of vertical inheritance as described previously (12, 20, 63, 64). Conversely, we observed the great diversity of host types in some subgroups, which could be partly explained by the ability of HLEs to undergo horizontal transfer (18, 65). Many domains were found to be limited within specific subgroups, further supporting the classification of subgroups and suggesting their potential as phylogenetic markers.

Some groups seemed to be devoid of HLEs, e.g., no HLE was detected among 29 sampled distinct birds. The mechanisms explaining this observation appear to be worth exploring in future research.

Furthermore, in the *A. mexicanum* giant genome (28.2 Gbp), we observed a remarkable divergence of *Helentrons* that formed the subgroup d, indicating the success of this subgroup in this species. However, in the larger giant lungfish *N. forsteri* genome (34.6 Gbp), we did not observe a comparable divergence of *Helentrons*. Future investigations on HLEs in giant genomes could be done to analyse how HLE evolved in these genomic landscapes.

In conclusion, this work provides a comprehensive and robust solution for improving HLE annotations in genomes. In particular, HELIANO’s ability to generate a novel annotation on full-length HLEs from a large set of samples makes it a unique and valuable tool for the scientific community.

## Data and Resource availability

Data and tools to prepare all supplementary data are available at https://zenodo.org/records/10625090. The source code of HELIANO is available from https://zenodo.org/records/10625240 and on github (https://github.com/Zhenlisme/heliano/).

## Authors contributions

Conceptualization: ZL CG HP NP; Data Curation: ZL NP; Funding Acquisition: ZL NP; Investigation: ZL ; Methodology: ZL HP; Project Administration: CG HP NP ; Resources: HP NP; Software: ZL ;Supervision: CG NP; Validation: ZL CG HP NP; Writing – Original Draft: ZL ;Writing – Review & Editing: ZL CG HP NP.

