## Supplementary_Figures for "Systematic annotation of *Helitron*-like elements in eukaryote genomes using HELIANO"

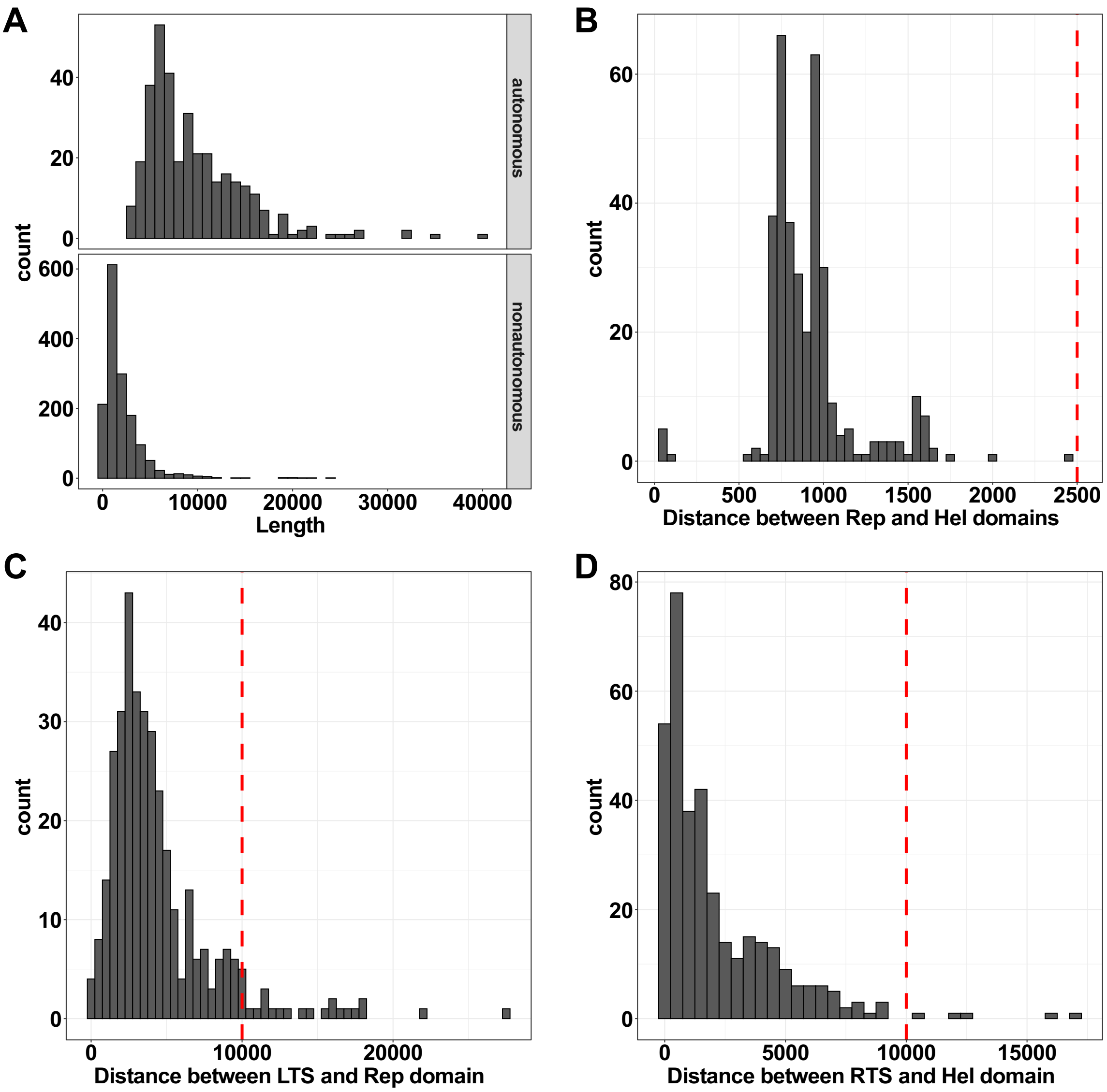


**Supplementary Figure 1.** Features of HLEs collected from Repbase. (A) Distribution of HLEs length. (B) Distriution of the distance between the 3' end of REP domain and 5' end of Helicase domain of HLEs. (C) Distribution of the distance between 5' end of LTS and 5' end of REP domain of HLEs. (D) Distribution of the distance between 3' end of Helicase domain and 3' end of RTS of HLEs. Red vertical lines indicate values used as default parameters in HELIANO. Length are expressed in bp.

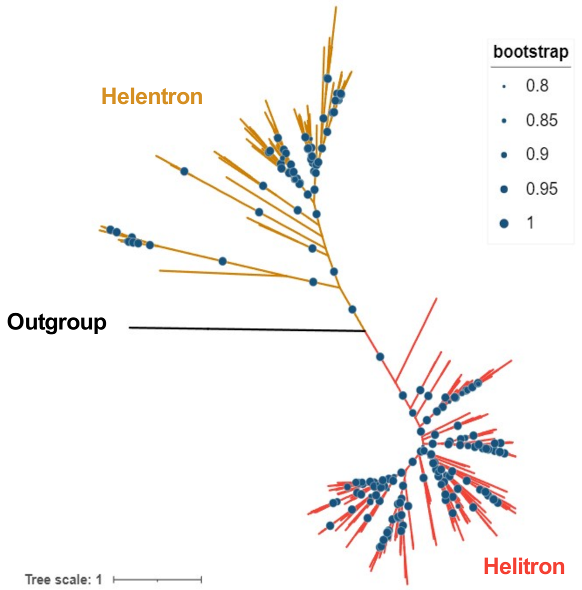


**Supplementary Figure 2**. Phylogenetic relationships of HLEs Rep domains. The clade highlighted in red encompass *Helitron* variants and the clade highlighted in orange encompass *Helentron* variants. The rep catalytic protein sequence of geminivirus (WP_015060107.1) was set as outgroup. This maximum likelihood estimation tree of Rep domains has a LogLk = -66062.678. Only bootstrap value greater than 0.8 are shown.

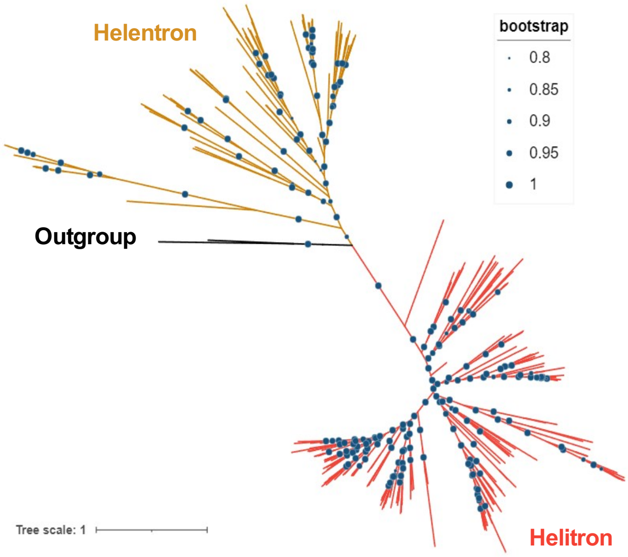


**Supplementary Figure 3**. Phylogenetic relationships of HLEs helicase domains. The clade highlighted in red encompass *Helitron* variants and the clade highlighted in orange encompass *Helentron* variants. The protein sequences of *Myroides phaeus* Helicase (WP_090404604.1) and of *Candidatus Collierbacteria* PIF1 helicase (KKT34677.1) were set as outgroups. This maximum likelihood estimation tree of Helicase domains of HLEs from Repbse has a LogLk = -108550.922. Only bootstrap value greater than 0.8 are shown.

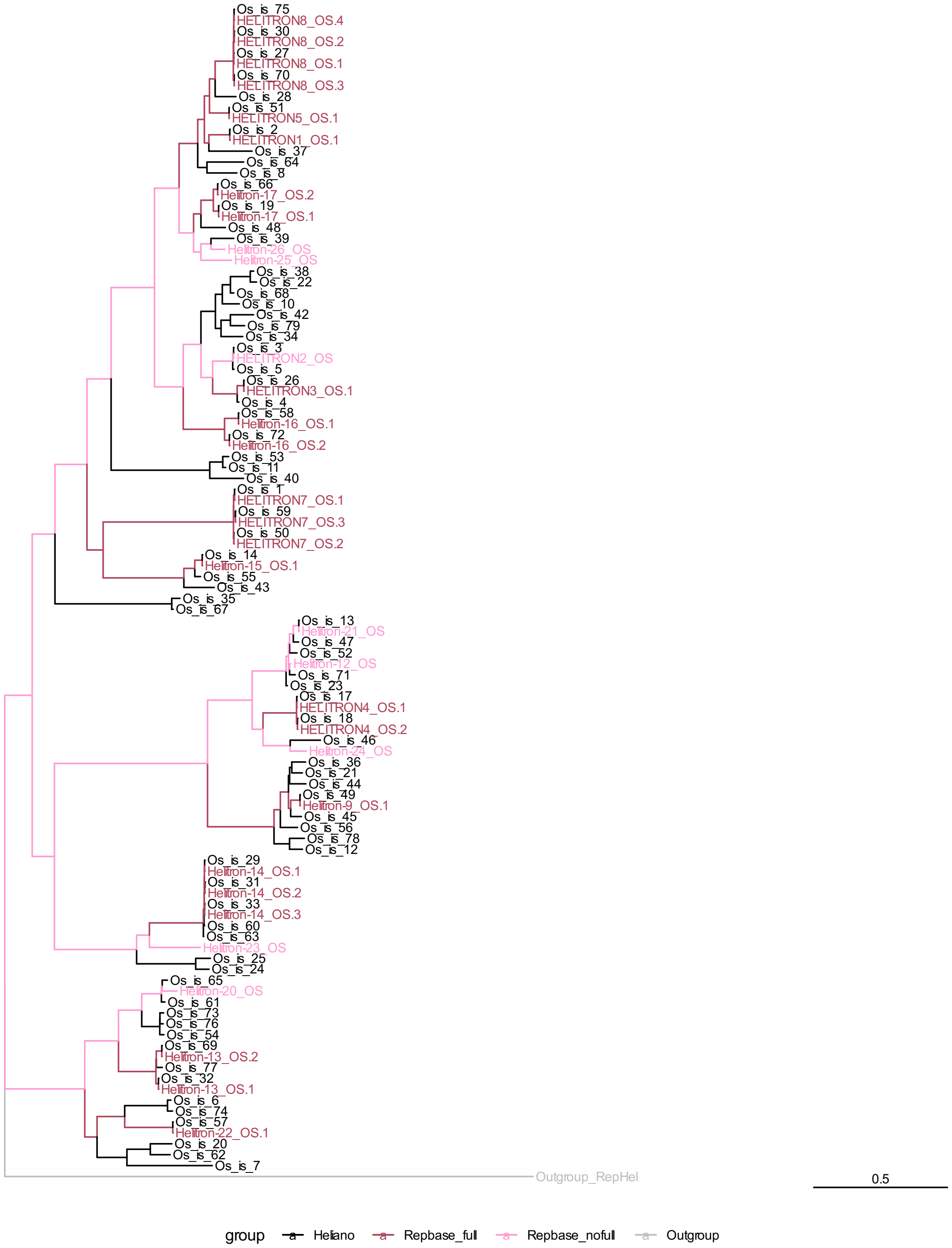


**Supplementary Figure 4**. Maximum likelihood estimation tree of transposases of *Helitron* insertions in *Oryza sativa* genome (LogLk = - 64380.460). Sequences labelled in red indicate transposases that were extracted from Repbase and present in *O. sativa* genome as complete copies. Sequences labelled in pink indicate transposases that were extracted from Repbase and absent in *O. sativa* genome as complete copies. Sequences labelled in black indicate transposases extracted from HELIANO annotation. The outgroup (Outgroup_RepHel in grey) was created by concatenating geminivirus rep catalytic protein and helicase protein of *Myroides phaeus*.

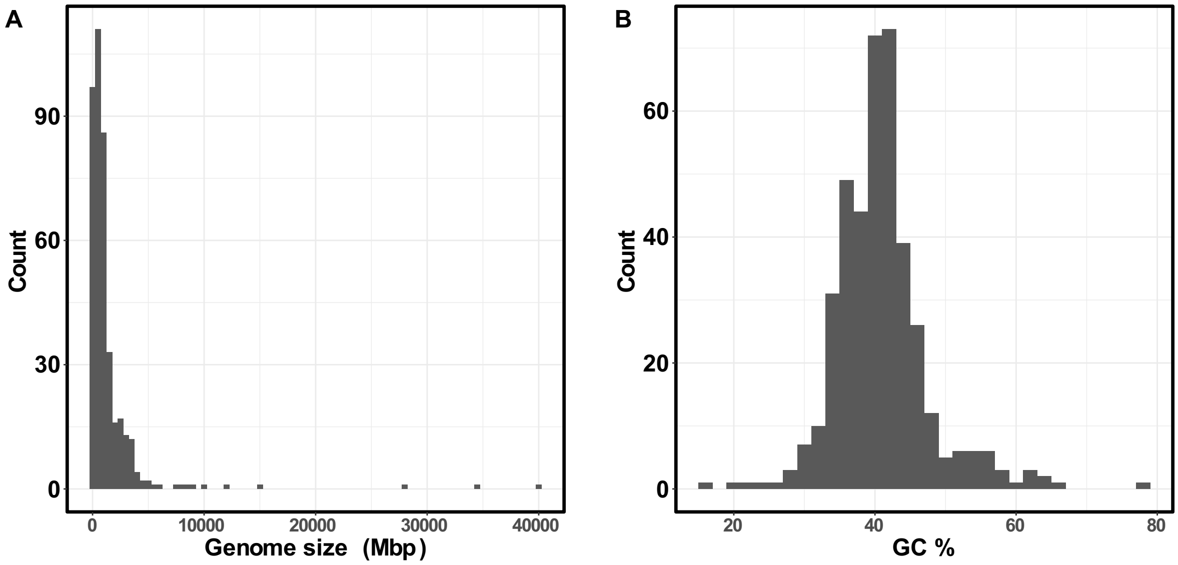


**Supplementary Figure 5**. Variation of genome size (A) and GC content (B) of 404 sampled genomes.

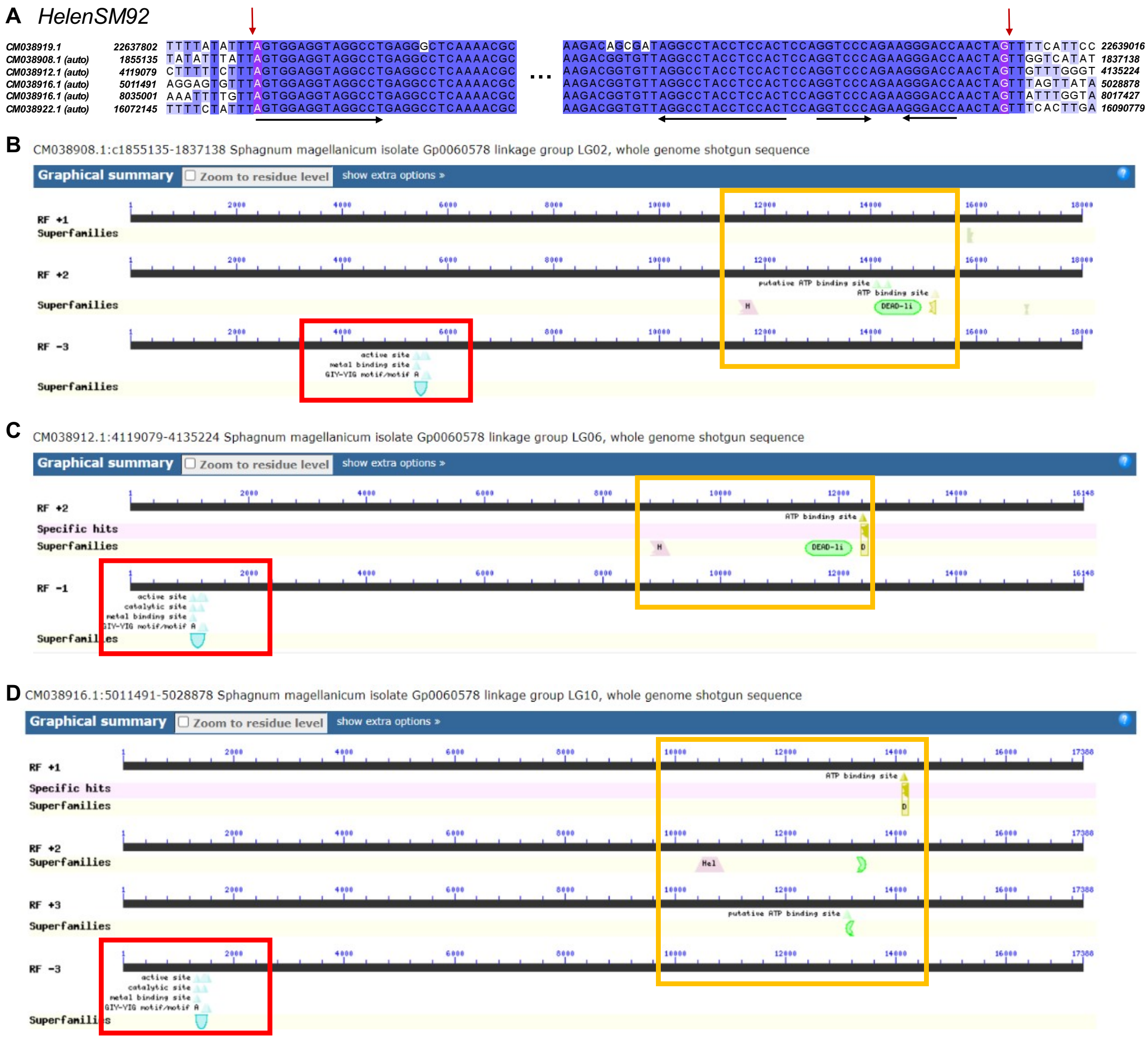


**Supplementary Figure 6**. Example of *Helentrons* in land plants. (A) Multiple alignments of selected *Helentron* insertions of HelenSM92 family in *Sphagnum magellanicum* genome. (B, C, D) Schemes showing the conserved domains for three *Helentron* insertion of HelenSM92 family. The GIY-YIG domains are highlighted in red rectangles and the HLE Rep-Hel domains are highlighted in orange rectangles. Image from the Conserved Domain Database (CDD) search tool (Lu et al. 2020).

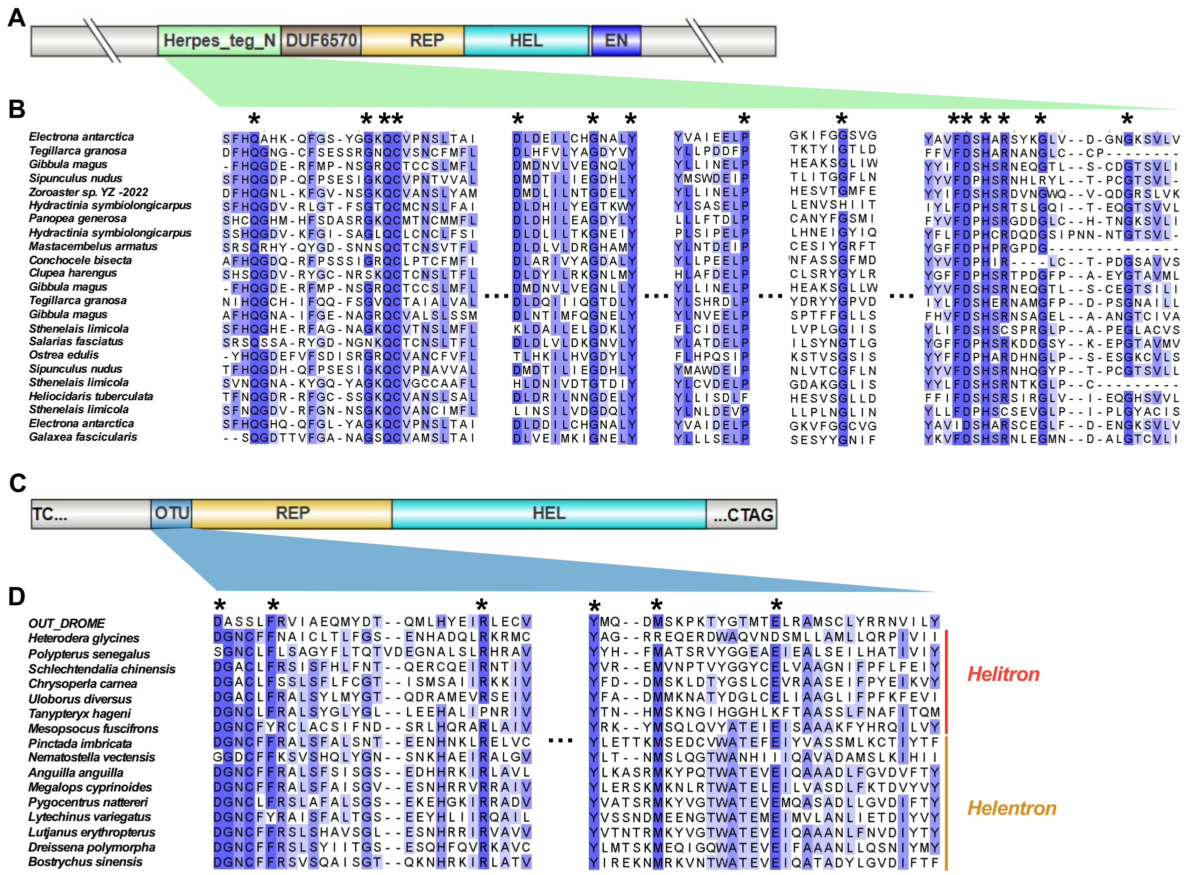


**Supplementary Figure 7**. Domains organisation of canonical *Helentrons* Rep/Hel proteins. (A) Domain organization of an *Helentron* Rep/Hel protein containing Herpes_teg_N, DUF6570 *and* EN domain. (B) Multiple alignment of Herpes_teg_N domain sequence of *Helentrons* from different species. (C) Domain organization of an *Helentron* Rep/Hel protein containing an OTU domain. (D) Multiple alignment of OTU sequences of *Helitrons* and *Helentrons* from different species. Asterisk symbol indicates conserved motifs.

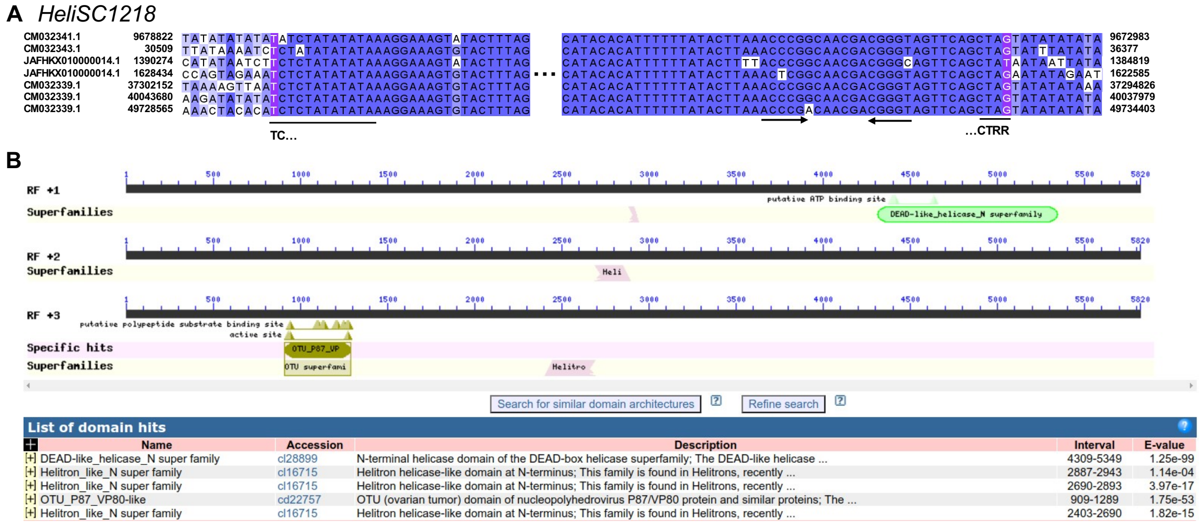


**Supplementary Figure 8**. The OTU domain in *Helitrons* of *Schlechtendalia chinensis*. (A) Multiple alignments of selected *Helitron* insertions of HeliSC1218 family. (B) conserved domains for one *Helitron* insertion of family HeliSC1218 (CM032341.1:c9678812-9672993). Image from the Conserved Domain Database (CDD) search tool (Lu et al. 2020).

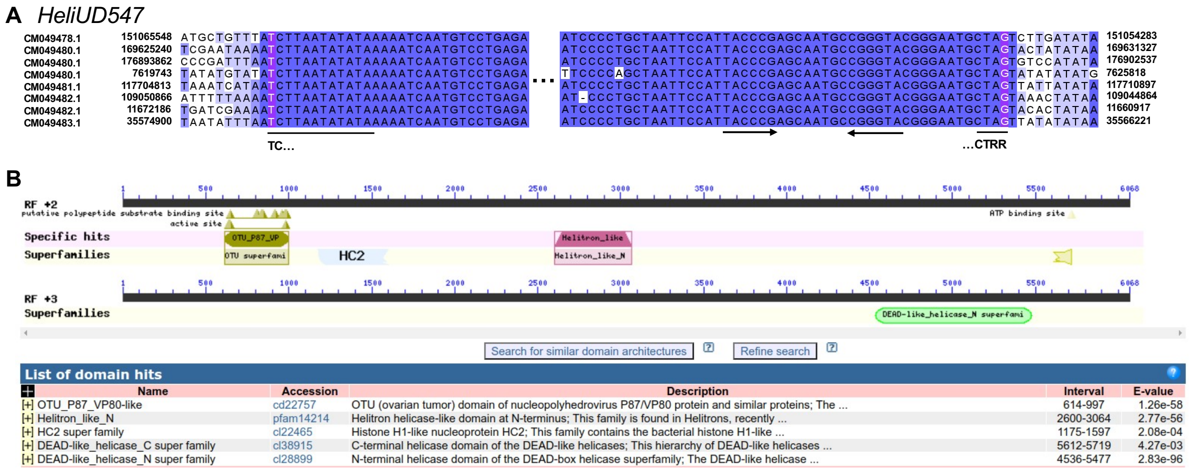


**Supplementary Figure 9**. The OTU domain in *Helitrons* of *Uloborus diversus*. (A) Multiple alignments of selected *Helitron* insertions of HeliUD547 family. (B) conserved domains for one *Helitron* insertion of family HeliUD547 (CM049480.1:169625250-169631317). Image from the Conserved Domain Database (CDD) search tool (Lu et al. 2020).

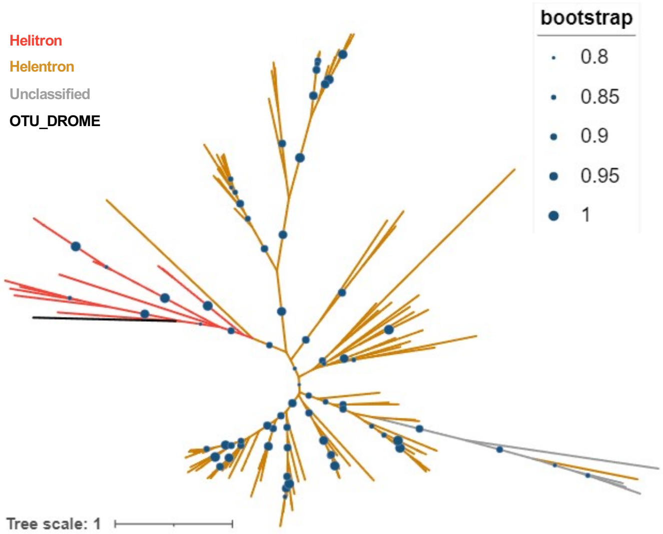


**Supplementary Figure 10**. Phylogeny of the OTU domain across HLEs. *Helitron* variant clade sequences are highlighted in red, *Helentron* variants are highlighted in orange and unclassified HLEs are in grey resembled. OTU_DROME protein (UniProtKB ID: P10383) was used as outgroup. Only bootstrap value greater than 0.8 are shown in this Maximum likelihood estimation tree.

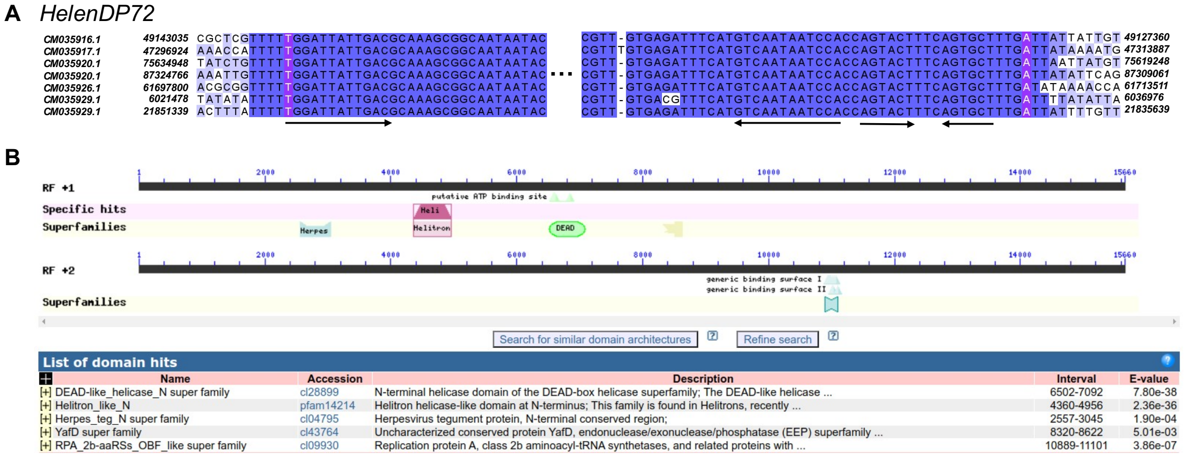


**Supplementary Figure 11**. The Herpes_teg_N domain in an *Helentron* of *Dreissena polymorpha*. (A) Multiple alignments of selected *Helentron* insertions of HelenUD547 family. (B) Conserved domains for one *Helentron* insertion of HelenUD547 family (CM035916.1:c49143025-49127369). Image from the Conserved Domain Database (CDD) search tool (Lu et al. 2020).

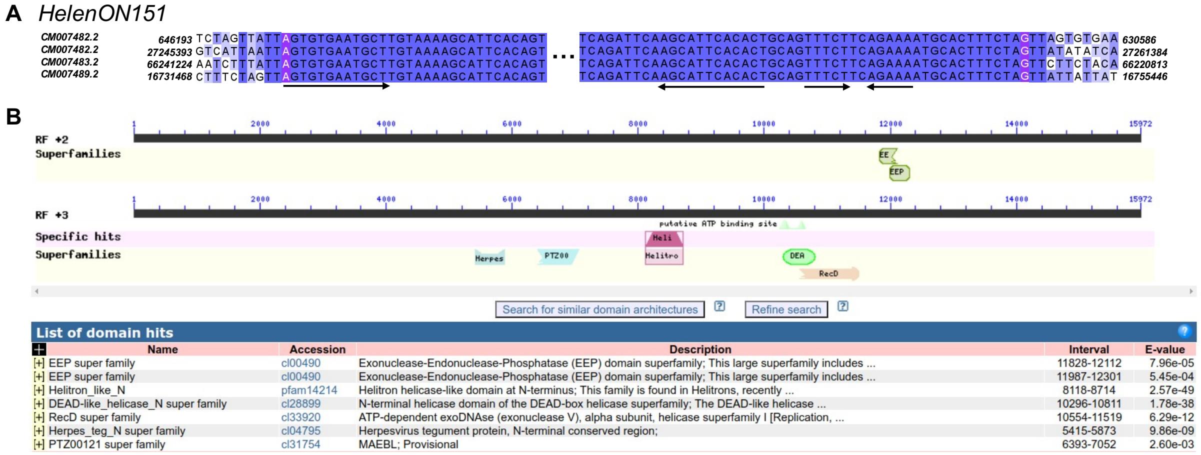


**Supplementary Figure 12**. The Herpes_teg_N domain in *Helentron* of *Oreochromis niloticus*. (A) Multiple alignments of selected *Helentron* insertions of HelenON151 family. (B) Conserved domains of one *Helentron* insertion of HelenON151 family (CM007482.2:27245403-27261371). Image from the Conserved Domain Database (CDD) search tool (Lu et al. 2020).

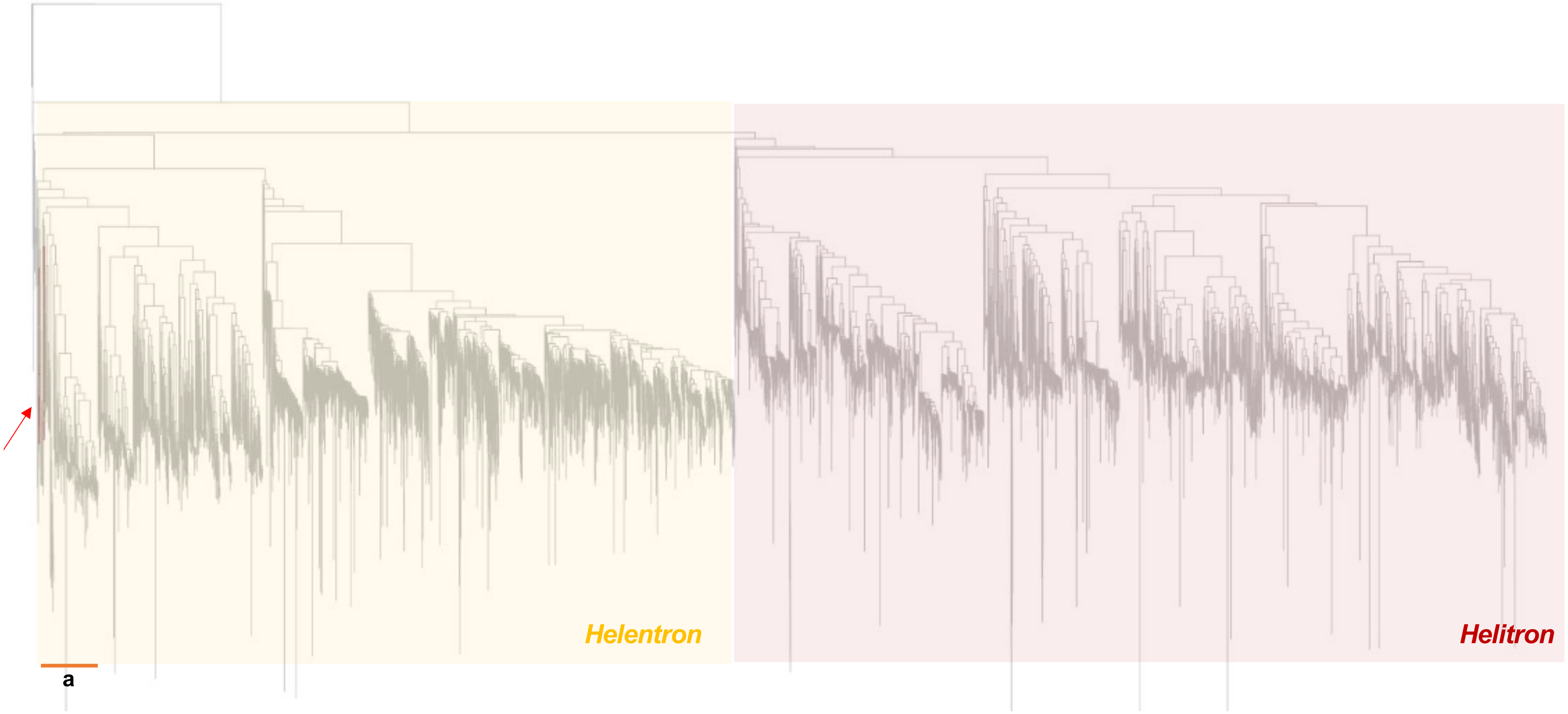


**Supplementary Figure 13**. Phylogenetic relationships of previously identified *Helitron2* sequences. The FoHeli2, FoHeli1, FOYG_17266, FOYG_17354 sequences are highlighted in red and pointed by a red arrow (Chellapan et al. 2016).

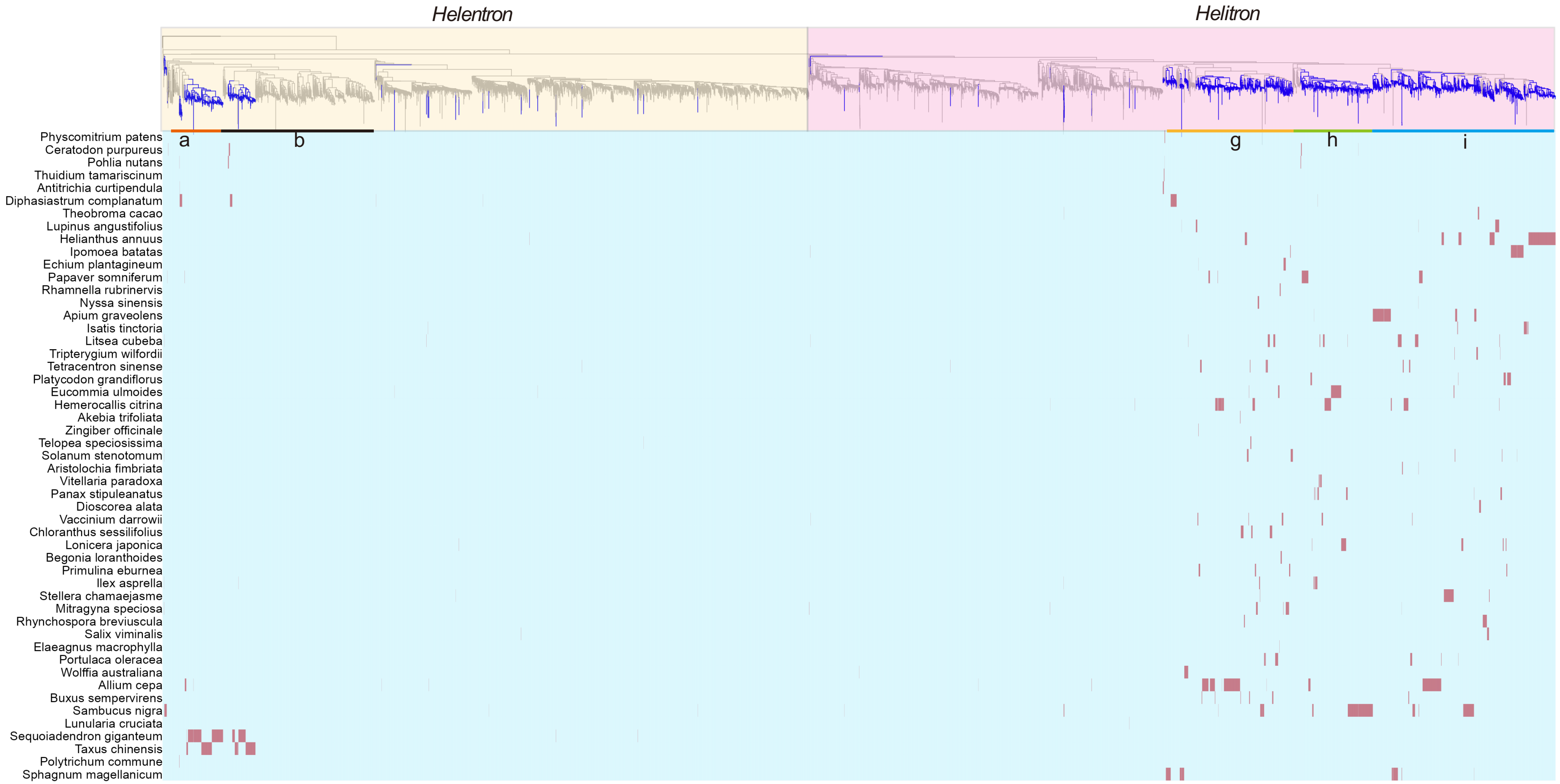


**Supplementary Figure 14**. Phylogenetic analysis of HLE sequences from land plant genomes. The branches highlighted in blue indicated HLE sequences from land plant genomes. In the bottom heat plot, the presence of HLE is highlighted in red.
